# iRQC, a surveillance pathway for 40S ribosomal quality control during mRNA translation initiation

**DOI:** 10.1101/2021.04.20.440649

**Authors:** Danielle M. Garshott, Heeseon An, Elayanambi Sundaramoorthy, Marilyn Leonard, Alison Vicary, J. Wade Harper, Eric J. Bennett

**Affiliations:** Section of Cell and Developmental Biology, Division of Biological Sciences; University of California-San Diego, La Jolla, CA, 92093; USA; Department of Cell Biology, Blavatnik Institute, Harvard Medical School; Boston, MA, 02115; USA

## Abstract

Since multiple ribosomes can engage a single mRNA, nonuniform ribosome progression can result in collisions. Ribosome collisions during translation elongation elicit a multifaceted ribosome-associated quality control (RQC) response. Despite advanced mechanistic understanding of translation initiation, a parallel RQC pathway that acts on collided preinitiation complexes has not been described. Here, we show that blocking progression of scanning or elongating ribosomes past the start codon triggers uS3 and uS5 ribosomal ubiquitylation. We demonstrate that conditions that activate the integrated stress response can also induce preinitiation complex collisions. The ubiquitin ligase, RNF10, and the deubiquitylating enzyme, USP10, are the key regulators of uS3 and uS5 ubiquitylation. Prolonged uS3 and uS5 ubiquitylation results in 40S, but not 60S, ribosomal protein degradation in an autophagy-independent manner. This study identifies a distinct arm in the RQC pathway, initiation RQC (iRQC), that acts on pervasive ribosome collisions during translation initiation to modulate translation activity and capacity.

## Introduction

Translation is the critical process that decodes the genetic blueprint into functional proteins. The three canonical steps of translation; initiation, elongation, and termination, are each highly regulated in order to tune protein biogenesis to match metabolic demands (Hershey et al., 2019). Of these, the mechanisms that regulate translation initiation are the most well-characterized (Hinnebusch, 2014; Pelletier and Sonenberg, 2019). Translation initiation involves assembly of the 43S preinitiation complex, comprised of the 40S ribosomal subunit, ternary complex (TC; eIF2, GTP, Met-tRNAi) and multiple initiation factors (Hinnebusch, 2017). The 43S is then recruited to the mRNA by the cap binding complex (eIF4F) and begins scanning in the 5’ to 3’ direction. Upon start codon recognition, the 60S ribosomal subunit joins and elongation ensues (Jackson et al., 2010). Translation is a resource intensive process, and diverse protein homeostasis stressors activate pathways that attenuate translation initiation to repress translational output (Leibovitch and Topisirovic, 2018). One such pathway is the integrated stress response (ISR). Varied cellular stimuli including viral infection and unfolded protein accumulation within the endoplasmic reticulum trigger the ISR which results in phosphorylation of the alpha subunit of the trimeric initiation factor eIF2 and subsequent translation initiation attenuation. (Pakos-Zebrucka et al., 2016). Both a failure to activate ISR and persistent ISR activation have physiological consequences, underscoring the importance of dynamic translation regulatory pathways (Back et al., 2009; Costa-Mattioli and Walter, 2020). While the ISR and pathways that downregulate translation initiation are highly dynamic, our current understanding is limited to how these responses facilitate short term changes in translational capacity. How translational capacity is rewired to reflect long-term changes in cell proliferation and metabolic demand remains unclear.

While most translation events terminate in successful protein biogenesis, cis-acting features of the mRNA or nascent chain can result in abortive translation termination (Hinnebusch et al., 2016). Defects in either the emerging nascent polypeptide or translating mRNA can cause ribosomes to experience prolonged stalls during elongation which can subsequently result in 80S ribosome collisions and elicit a multifaceted ribosome-associated quality control (RQC) pathway (Inada, 2020; Joazeiro, 2019). Components of the RQC act to degrade the truncated nascent chain, destroy the associated mRNA, and recycle the ribosomal subunits (Joazeiro, 2019; Meydan and Guydosh, 2020). Current models suggest that ribosome collisions are the integral first signal necessary to recruit factors that facilitate downstream RQC activities (D’Orazio and Green, 2021; Ikeuchi et al., 2019; Juszkiewicz et al., 2018; Simms et al., 2017). Protein ubiquitylation plays two critical roles during RQC. The first involves conserved regulatory ribosomal ubiquitylation (RRub) of 40S proteins eS10 (RPS10) and uS10 (RPS20) mediated by the E3 ligase ZNF598 (Juszkiewicz and Hegde, 2017; Matsuo et al., 2017; Sundaramoorthy et al., 2017). The second involves an additional ligase, Listerin, which is recruited to the 60S subunit, post-80S ribosome splitting, to catalyze nascent polypeptide chain ubiquitylation and subsequent degradation (Shao and Hegde, 2014; Shao et al., 2013). While these ubiquitylation events are well characterized, additional sites of ribosome ubiquitylation have been described that either play a direct role within the RQC (Ikeuchi et al., 2019; Saito et al., 2015) or operate outside of the RQC and have uncharacterized roles (Higgins et al., 2015; Montellese et al., 2020; Silva et al., 2015), suggesting ribosome ubiquitylation may regulate multiple steps during translation.

Dynamic feedback regulation between elongation and initiation meters ribosome traffic along mRNAs. Elevation in RQC activity due to increases in elongating ribosome collisions can indicate an overabundance of ribosome density on transcripts. Compensatory decreases in translation initiation rates can reduce ribosome collisions and RQC activity (Juszkiewicz et al., 2018). Further, recent studies have defined a collision-induced feedback loop that downregulates translation initiation. Following ribosome collisions, in a ZNF598-independent manner, the collision-sensor EDF1 recruits the translation repressors GIGYF2 and 4EHP to inhibit translation of stall-inducing transcripts (Juszkiewicz et al., 2020; Sinha et al., 2020). A separate study also demonstrated that the same translation repressors, GIGYF2 and 4EHP, when deleted, increased translation of a stall-inducing reporter (Hickey et al., 2020). These studies highlight the requirement for dynamic coordination between elongation and initiation rates to regulate elongation collision frequency. While elongating ribosome collisions and the corresponding RQC pathway have been well-established, a similar quality control pathway that operates within the initiation phase of translation has not been described.

Here we establish a new surveillance pathway, iRQC, in which regulatory ribosomal ubiquitylation demarcates collided preinitiation complexes. We demonstrate that impeding progression of scanning or elongating ribosomes near start codons induces site-specific uS3 (RPS3) and uS5 (RPS2) ribosome ubiquitylation. We identify and characterize the ubiquitin ligase RNF10, and the deubiquitylating enzyme, USP10 as the key ubiquitin pathway enzymes that regulate iRQC-specific ribosomal ubiquitylation. Loss of USP10 function resulted in enhanced uS5 and uS3 ubiquitylation in the absence of exogenous stressors indicating that preinitiation collisions are pervasive. We show that prolonged uS3 and uS5 ubiquitylation induces selective degradation of 40S ribosomal proteins in an autophagy-independent manner. Our results establish parallel but distinct RQC pathways that act on ribosome collisions during the elongation (eRQC) or initiation (iRQC) phases of translation. As iRQC pathway activation results in 40S ribosomal protein degradation, the frequency of initiation collisions can be used as a rheostat to balance ribosome abundance and overall translational capacity with metabolic demands.

## Results

### Translation initiation inhibition triggers 40S ribosomal ubiquitylation

We had previously demonstrated that translation inhibition mediated by Harringtonine, which blocks progression of 80S ribosomes at the start codon without impacting elongating or scanning ribosomes (Fresno et al., 1977), induces site-specific ubiquitylation events on the 40S ribosomal proteins uS5 (RPS2) and uS3 (RPS3)(Higgins et al., 2015). Intriguingly, the uS5 and uS3 ubiquitylation sites are similarly positioned within the ribosome as the uS10 (RPS20) and eS10 (RPS10) ubiquitylation sites that are required for RQC events during elongation collisions (Figure 1A) (Ikeuchi et al., 2019; Juszkiewicz et al., 2018; Juszkiewicz and Hegde, 2017; Matsuo et al., 2017; Sundaramoorthy et al., 2017). Since uS3 and uS5 ubiquitylation do not function within the characterized RQC pathway (Garshott et al., 2020), we hypothesized that preinitiation complex collisions during the mRNA scanning phase of translation initiation trigger uS3 and uS5 ubiquitylation. If uS3 and uS5 ubiquitylation occurs in response to collided preinitiation complexes, then impeding progression of scanning or elongating ribosomes near start codons should induce ribosome ubiquitylation. To test this idea, we first treated cells with increasing concentrations of harringtonine (HTN). Consistent with previous results, HTN treatment resulted in rapid and robust uS5 and uS3 ubiquitylation (Figures 1B and 1C)(Higgins et al., 2015). Treatment with either HTN or lactimidomycin (LTM), a functionally similar but mechanistically distinct compound (Lee et al., 2012; Schneider-Poetsch et al., 2010), induced uS5 and uS3 ubiquitylation that was detectable after 5 minutes and further increased over time (Figure 1D). Because uS10 and eS10 ubiquitylation occurs upon 80S ribosome collisions during elongation, the observed HTN-induced uS3 and uS5 ubiquitylation may occur when scanning 43S ribosomes collide with stalled 80S ribosomes poised at the initiation codon.

**Figure 1.**
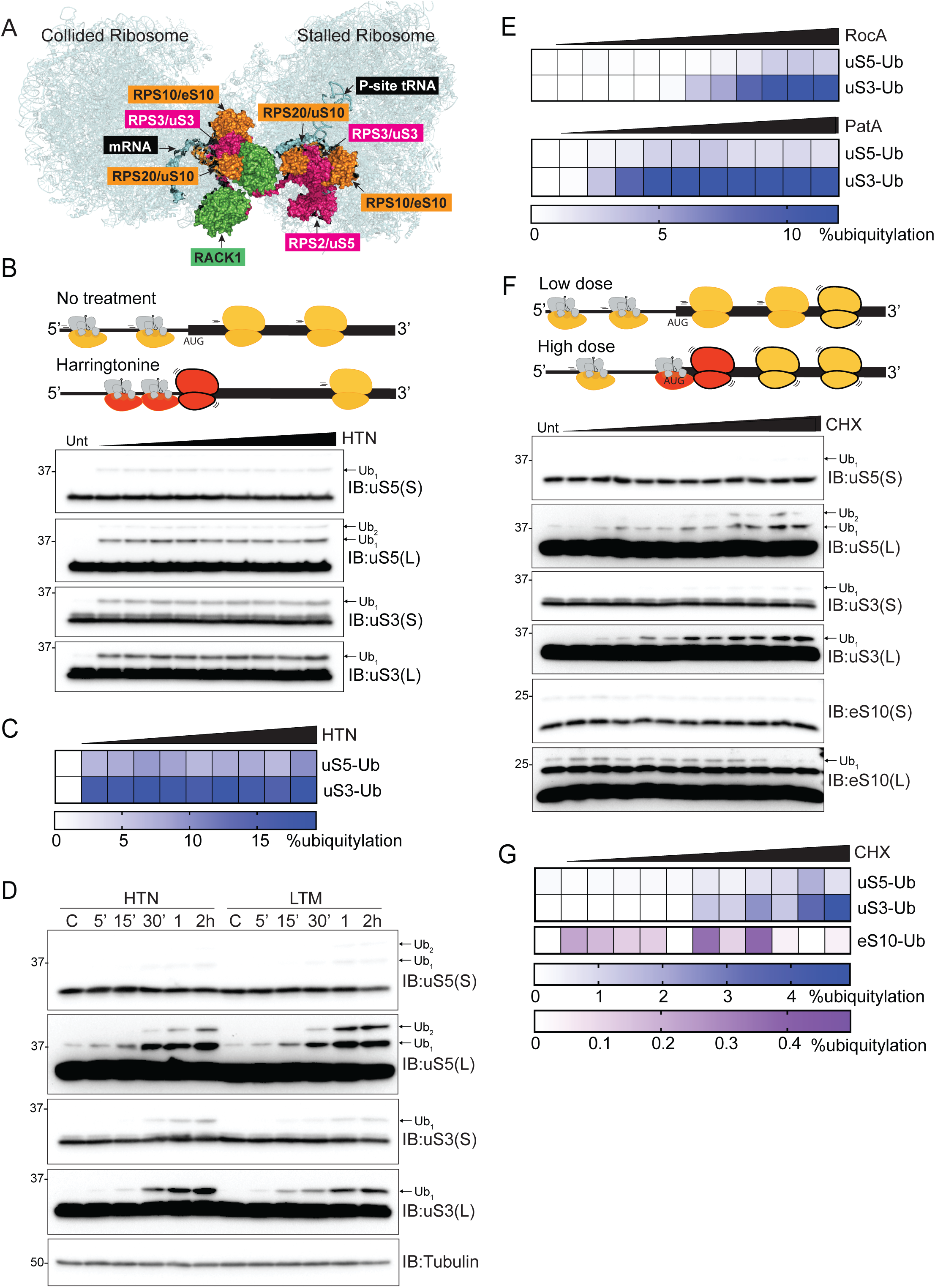
Translational initiation inhibition induces ribosomal ubiquitylation, See also Figure S1. (A) Collided disome structure with ubiquitylated 40S proteins that function either within (uS10/RPS20, eS10/RP10 shown in orange) or outside (uS3/RPS3, uS5/RPS2 shown in magenta) the characterized RQC pathway. RACK1 (green) mRNA (blue), and P-site tRNA (blue) are indicated. Structure from PDB:6HCQ and 6HCM. (B) (top) Schematic of Harringtonine (HTN)-induced translation inhibition. Preinitiation complexes are shown scanning in the 5’UTR. Collided ribosomes are shown in red and HTN-inhibited ribosomes are indicated by black outline. (bottom) Cell extracts from 293T cells treated with increasing doses of HTN (0.039-10ug/ml) for two hours were analyzed by SDS-PAGE and immunoblotted (IB) with the indicated antibodies. For all blots, the ubiquitin-modified ribosomal protein is indicated by the arrow. S and L denote short and long exposures, respectively. (C) Percent ubiquitylated uS3 and uS5 quantified from immunoblots in B. (D) Cell extracts from 293T cells treated with harringtonine (HTN) (2ug/ml) or lactimidomycin (LTM) (5uM) for the indicated times were analyzed by SDS-PAGE and immunoblotted with the indicated antibodies. (E) Quantification of uS3 or uS5 percent ubiquitylation from 293T cells treated with increasing doses of either rocaglates (RocA) (0.031-3.2uM) or patamineA (PatA) (0.31-100nM) from blots in S1A,B. (F) (top) Schematic of the differential impact on initiation collisions upon treatment with high dose versus low dose elongation (cycloheximide) inhibitors. Collided ribosomes are shown in red and CHX-inhibited ribosomes are indicated by black outline. (bottom) Cell extracts from 293T cells treated with increasing concentration of cycloheximide (CHX) (0.01-500ug/ml) for two hours were analyzed by SDS-PAGE and immunoblotted with the indicated antibodies. (G) Quantitative representation of uS3, uS5, and eS10 percent ubiquitylation from F.

We then utilized characterized inhibitors that impede the mRNA scanning step of translation initiation to examine if inhibiting progression of ribosomes past the start codon induces ribosome ubiquitylation. Addition of rocaglates (RocA)(Iwasaki et al., 2016) or pateamine A (PatA)(Low et al., 2005), which inhibit the RNA helicase eIF4A and impair mRNA scanning, induced uS5 and uS3 ubiquitylation in a dose dependent manner (Figures 1E and S1A-S1B). As HTN, LTM, RocA, and PatA each are expected to trigger collisions between either multiple 43S preinitiation complexes scanning within the 5’UTR or between scanning preinitiation complexes and a stalled 80S, these results suggest that collisions involving at least one preinitiation complex trigger uS5 and uS3 ubiquitylation.

Cycloheximide (CHX) and anisomycin (ANS), translation elongation inhibitors that act on elongating 80S ribosomes, induce both RQC-required uS10 and eS10 ubiquitylation and RQC-independent uS3 and uS5 ubiquitylation (Higgins et al., 2015; Juszkiewicz and Hegde, 2017; Sundaramoorthy et al., 2017). The demonstration that maximal elongation collisions and uS10 and eS10 ubiquitylation occur with low dose, rather than high dose treatment of elongation inhibitors was a critical result establishing that ribosome collisions are the key event leading to ribosomal ubiquitylation and RQC pathway activation (Juszkiewicz et al., 2018; Simms et al., 2017). In contrast to what was observed for elongation collisions, we predicted that uS3 and uS5 ubiquitylation would be optimal at high concentrations of elongation inhibitors, as collisions with scanning ribosomes would result only if enough 80S ribosomes accumulated in proximity to the start codon (Figure 1F). Treating cells with increasing CHX concentrations confirmed this prediction (Figures 1F and 1G). The ubiquitylation of uS5 and uS3 increased with elevating CHX concentrations, as opposed to eS10 ubiquitylation, which was induced at low CHX concentrations and diminished at high concentrations (Figures 1F and 1G). These results establish that preinitiation complex collisions induce uS3 and uS5 ubiquitylation.

### ISR activation elicits preinitiation collisions

We previously demonstrated that uS3 and uS5 ubiquitylation occurs upon activation of the integrated stress response (ISR) in an eIF2*α* phosphorylation-dependent manner (Higgins et al., 2015). ISR stimulated eIF2*α* phosphorylation inhibits translation initiation through depletion of the ternary complex, which consists of methionyl-initiator tRNA (Met-tRNAi) and guanosine triphosphate (GTP)–bound eIF2 (Costa-Mattioli and Walter, 2020; Hinnebusch, 2014). It was initially puzzling why stressors that reduce translation initiation activity would result in uS3 and uS5 ubiquitylation if those modifications demarcate preinitiation collisions. One explanation would be that ISR activation also induces preinitiation collisions. We noticed that distinct ISR agonists increased uS3 and uS5 ubiquitylation to varying degrees, with those inducing low levels of eIF2*α* phosphorylation resulting in higher uS3 and uS5 ubiquitylation (Figures 2A and S1C). Notably, high concentration sodium arsenite (NaAsO2) treatment resulted in the greatest extent of eIF2*α* phosphorylation, but poorly stimulated uS3 or uS5 ubiquitylation (Figures 2A and S1C).

**Figure 2.**
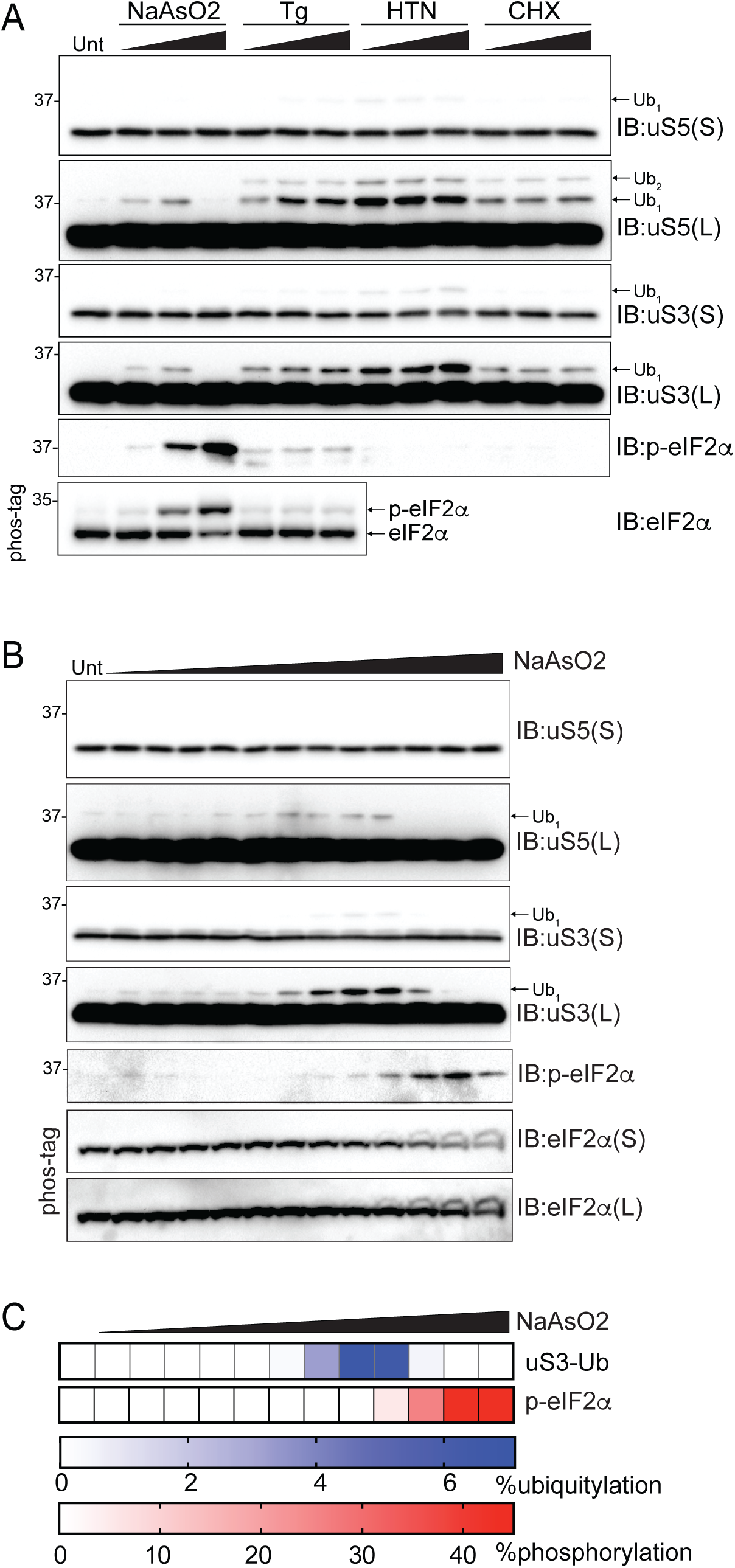
Integrated stress response activation results in preinitiation complex collisions, See also Figure S1. (A) Cell extracts from 293T cells treated with increasing concentrations of thapsigargin (Tg) (0.1, 1, 5uM), HTN (0.25, 2, 10ug/ml), CHX (5, 50, 150ug/ml), or sodium arsenite (NaAsO2) (10, 125, 1mM) for 2 hours were analyzed by SDS-PAGE and immunoblotted with the indicated antibodies. For all blots, the ubiquitin-modified ribosomal protein is indicated by the arrow. S and L denote short and long exposures, respectively. (B) Cell extracts from 293T cells treated with increasing concentration of NaAsO2 (488nM-1mM) for two hours were analyzed by SDS-PAGE and immunoblotted with the indicated antibodies. (C) Quantification of uS3 percent ubiquitylation and eIF2*α* percent phosphorylation (from Phos-tag gels) following NaAsO2 treatment from B.

We reasoned that high stoichiometry eIF2*α* phosphorylation would inhibit ribosome collisions during initiation by completely depleting GTP-bound ternary complexes and blocking translation initiation. In contrast, ISR agonists that induce low levels of eIF2*α* phosphorylation may allow for distinct scanning preinitiation complexes with either GDP or GTP-bound ternary complexes on a single mRNA. Scanning complexes with or without GTP differentially recognize suboptimal start codons (Hinnebusch, 2017). Thus, scanning ribosomes with GDP-bound ternary complex may progress past some start codons, while other scanning ribosomes containing GTP, on the same mRNA, will transition to elongating ribosomes. We hypothesized that these mixed populations of scanning and elongating ribosomes within the coding sequence stimulate collisions. To test if low stoichiometry eIF2*α* phosphorylation induces preinitiation collisions and subsequent ribosome ubiquitylation, we treated cells with a range of sodium arsenite concentrations and quantified uS3 and uS5 ubiquitylation and eIF2*α* phosphorylation. Arsenite concentrations that induced less than 5% eIF2*α* phosphorylation resulted in maximal uS3 and uS5 ubiquitylation whereas conditions in which eIF2*α* was phosphorylated in excess of 40% did not induce ubiquitylation (Figure 2B and 2C). These results are consistent with our hypothesis that ISR activation resulting in low stoichiometry eIF2*α* phosphorylation allows for collisions between either multiple preinitiation complexes or preinitiation complexes and 80S elongating ribosomes. Our results demonstrate that distinct, conserved ribosomal ubiquitylation events operate within separate RQC pathways which we classify as elongation RQC (eRQC) and initiation RQC (iRQC).

### Ribosome profiling analysis of preinitiation collisions

We next sought to determine if uS3 and uS5 ubiquitylation is enriched within ribosomal populations that may contain collided translation preinitiation complexes. As the mRNA present within collided 80S ribosomes is resistant to RNase treatment due to the compact nature of the collided disome structure (Juszkiewicz et al., 2018), we initially examined ribosome protein abundance across sucrose gradients from lysates treated with RNaseA. We compared untreated and HTN treated cells and observed that HTN treatment resulted in a noticeable broadening of the canonical 80S monosome peak, with a skew toward the lower density fractions (Figure 3A). Immunoblotting revealed that ribosomes with maximal HTN-induced uS3 and uS5 ubiquitylation migrated within fraction 5 which is at the front edge of the traditional monosome peak (Figure 3B). Abundant ubiquitylation within the 40S peak, which may also contain individual 43S preinitiation complexes, was also observed. Interestingly, uS3 ubiquitylation was observed within the monosome peak in untreated cells, hinting that preinitiation collisions may occur in the absence of stress (Figure 3B).

**Figure 3.**
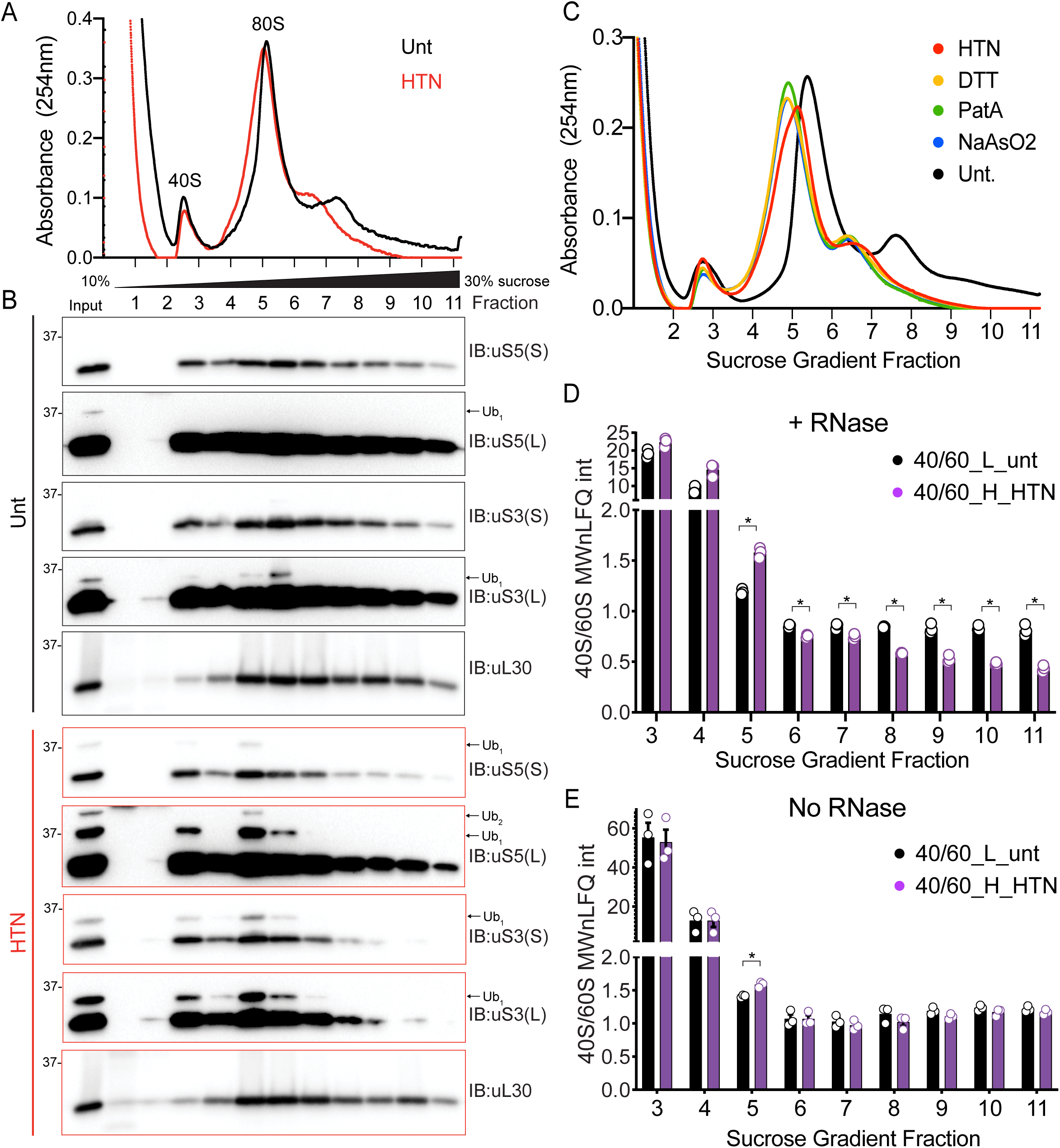
HTN induces 40S ubiquitylation in density gradient fractions with excess 40S relative to 60S ribosomal proteins, See also Figure S2. (A) RNaseA treated cell extracts from untreated (black line) or HTN (2ug/ml) treated (red line) 293T cells were fractionated on 10-30% sucrose gradients. The 254nm absorbance trace is depicted. (B) Fractions (designated in A) were analyzed by SDS-PAGE and immunoblotted with the indicated antibodies. The ubiquitin-modified ribosomal protein is indicated by the arrow. S and L denote short and long exposures respectively. (C) RNaseA treated cell lysates from untreated (black line), HTN (2ug/ml) (red line), DTT (5mM) (yellow line), PatA (100nM) (green line) or NaAsO2 (125uM) (blue line) treated 293T cells were fractionated on 10-30% sucrose gradients. The relative 254nm absorbance trace is depicted. (D,E) The ratio of the summed molecular weight (MW) normalized LFQ intensities 40S proteins:60S proteins from untreated, light labeled (black bars) or HTN treated, heavy labeled (purple bars) 293T cells. Cell extracts with either untreated (E) or treated (D) with RNaseA prior to density gradient centrifugation. Bars denote mean value for replicate experiments (n=3) with error bars displaying SEM. *=pvalue<0.05 by unpaired two-tailed Student’s t test.

In addition to HTN treatment, the widening and skewing of the monosome peak was observed in fractionated lysates from cells treated with DTT, PatA, or a moderate dose of NaAsO2, all of which induce uS5 and uS3 ubiquitylation (Figure 3C). While the monosome peak presumably contains predominately individual 80S complexes, this peak may also contain mRNAs with multiple loaded preinitiation complexes. According to this hypothesis, we should observe more 40S ribosome proteins relative to 60S proteins within the monosome peak under conditions that stimulate preinitiation collisions. We utilized SILAC-based quantitative proteomics to compare the abundance of 40S relative to 60S proteins across the sucrose gradient. Heavy-labeled HTN treated cells were mixed with untreated cells prior to lysis and density centrifugation. Directly comparing ribosome protein ratios revealed the expected increase in both 40S and 60S proteins in monosome-containing sucrose fractions in HTN-treated cells (Figures S2A-S2B; Table S1). We observed a significant increase in the 40S protein ratio compared to 60S in fraction 5, at the leading edge of the monosome peak, when comparing the summed and molecular weight normalized ion intensities from all 40S or 60S ribosomal proteins (Figure 3D; Table S1). This result is consistent with the hypothesis that HTN-induces preinitiation complex collisions which migrate within the canonical monosome fraction in sucrose gradients. RNase treatment also resulted in noticeable deviation from the expected 40S:60S ratio in polysome-containing fractions (Figure 3D). Because RNaseA-mediated rRNA degradation may be impacting the integrity of 40S or 60S subunits, we repeated the sucrose gradient analysis without RNase treatment. We observed robust uS3 ubiquitylation throughout the broad HTN-induced monosome peak that was absent in untreated samples (Figures S2C,D). Further, we observed an increase in 40S protein abundance relative to 60S only in fraction 5 from HTN-treated cells (Figures 3E and S2E,F, Table S1). While this analysis would benefit from an enhanced resolution of distinct ribosomal complexes that is not achievable with standard sucrose gradient centrifugation, these results suggest that HTN induces collisions between preinitiation complexes, which stimulates iRQC pathway activation.

### RNF10 catalyzes uS5 and uS3 ubiquitylation during iRQC

In order to determine the molecular consequence of initiation collisions and the role uS3 and uS5 ubiquitylation play during iRQC, we set out to identify the ubiquitin pathway enzymes that regulate uS3 and uS5 ubiquitylation. We utilized an siRNA-based loss-of-function screen targeting 18 known RNA-associated ubiquitin ligases and found that only depletion of RNF10 reproducibly prevented uS3 and uS5 ubiquitylation (Figures 4A and S3A-S3E). We then generated and verified RNF10 knockout cells using CRISPR-Cas9-based approaches and demonstrated that these cells completely lacked DTT-or ANS-induced uS3 and uS5 ubiquitylation (Figure 4B). To investigate the specificity of RNF10, we examined uS5, uS3, eS10, and uS10 ubiquitylation in 293 Flp-IN cells expressing inducible wild type RNF10. In the same manner that ZNF598 is specific in modifying eS10 and uS10 (Juszkiewicz and Hegde, 2017; Sundaramoorthy et al., 2017), RNF10 expression, in the absence of stress, resulted in enhanced uS3 and uS5 ubiquitylation but left eS10 and uS10 ubiquitylation largely unchanged (Figure 4C). Furthermore, in vitro ubiquitylation assays demonstrated that RNF10 maintains its ribosomal protein specificity when incubated with purified 40S subunits (Figures 4D and S3F). Collectively, these findings demonstrate that RNF10 is both necessary and sufficient to catalyze uS3 and uS5 ubiquitylation.

**Figure 4.**
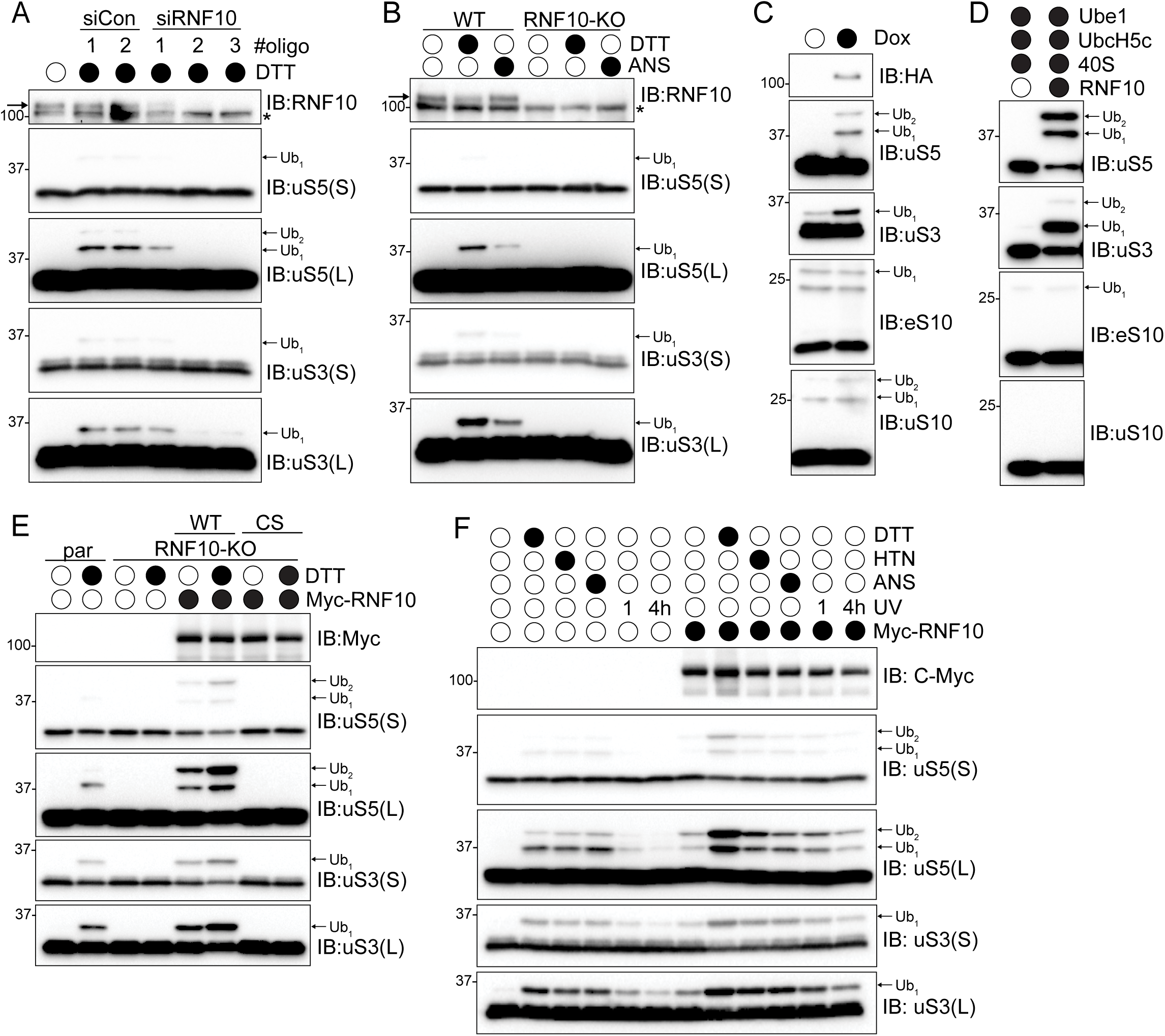
RNF10 catalyzes uS3 and uS5 ubiquitylation, See also Figure S3. (A) Cell lysates from 293T cells transfected with either control siRNA oligos or three separate siRNA oligos targeting RNF10, followed by treatment with dithiothreitol (DTT) for 2 hours were analyzed by SDS-PAGE and immunoblotted with the indicated antibodies. * indicates non-specific background signal. Arrow indicates RNF10-specific immunoreactivity. For all blots the ubiquitin-modified ribosomal protein is indicated by the arrow. S and L denote short and long exposures, respectively. (B) Cell extracts from parental 293T or RNF10 knockout (KO) cells were either untreated or treated with DTT (5mM) or anisomycin (ANS) (5ug/ml) then analyzed by SDS-PAGE and immunoblotted with the indicated antibodies. (C) HEK293-FlpIn cells expressing tet-inducible Flag-HA tagged RNF10 were treated with doxycycline (Dox) and cell lysates were analyzed by SDS-PAGE and immunoblotted with the indicated antibodies. (D) In vitro ubiquitylation assay utilizing purified 40S ribosomal subunits and RNF10. Samples were analyzed by SDS-PAGE and immunoblotted with the indicated antibodies. (E) RNF10 knockout (KO) cells were transfected with Myc-tagged wild type (WT) or inactive mutant (CS) RNF10 and parental 293T or RNF10-KO cells were either untreated or treated with DTT (5mM) for 2 hours. Cell lysates were analyzed by SDS-PAGE and immunoblotted with the indicated antibodies. (F) 293T cells with and without Myc-tagged wild type RNF10 expression were drug treated as indicated. UV indicates that cells were exposed to UV (0.02J/cm2) and were allowed to recover for 1 or 4 hours. Cell extracts were analyzed by SDS-PAGE and immunoblotted with the indicated antibodies.

What is the fate of the ubiquitylated 40S? To begin to address this question, we utilized an RNF10 overexpression system where we transiently overexpressed either wild type RNF10, or a catalytically inactive mutant (C225S) in RNF10 knockout (KO) cells to enhance uS5 and uS3 ubiquitylation both at basal conditions and upon conditions that stimulate preinitiation collisions. Expression of wild type, but not inactive RN10, rescued the ability to ubiquitylate uS3 and uS5 with or without DTT treatment (Figures 4E and S3G). Combining RNF10 overexpression with DTT treatment resulted in enhanced uS3 and uS5 ubiquitylation with more than 40% of total uS3 being ubiquitylated (Figures 4E and S3G). The extent of uS5 di-ubiquitin modification was also substantially increased upon RNF10 overexpression. Notably, total uS3 and uS5 protein abundance was also reduced upon RNF10 overexpression with and without DTT treatment (Figures 4E and S3G). This result suggests the RNF10-catalyzed 40S ubiquitylation acts to reduce 40S protein levels. Utilizing a panel of ISR and elongation inhibitors, we observed that RNF10 overexpression further heightened uS3 and uS5 ubiquitylation while having no additional effect on eS10 or uS10 modification (Figures 4F and S3H). Because RNF10 overexpression, in the absence of stressors, induced uS5 and uS3 ubiquitylation, we hypothesized that preinitiation complex collisions may occur under normal proliferative conditions and that the extent of ribosomal ubiquitylation may be limited by the cellular concentration of RNF10.

### USP10 antagonizes RNF10-dependent uS3 and uS5 ubiquitylation

The observed low levels of steady state uS3 and uS5 ubiquitylation suggested that either preinitiation collisions are rare events in the absence of stress, or that abundant deubiquitylating activity rapidly antagonizes RNF10-mediated ribosomal ubiquitylation. Our demonstration that RNF10 overexpression can stimulate uS3 and uS5 ubiquitylation at steady state argued that robust deubiquitylating activity antagonizes RNF10 ribosomal ubiquitylation. We serendipitously identified USP10 as the deubiquitylating enzyme responsible for removing ubiquitin from uS3 and uS5 while identifying and characterizing deubiquitylating enzymes that antagonize ZNF598 (Figure S4A) (Garshott et al., 2020). A recent study also identified USP10 as a ribosomal deubiquitylating enzyme (Meyer et al., 2020). Consistent with this previous work, USP10 knockout (KO) cells display constitutively high levels of not only ubiquitylated eS10 and uS10, but also ubiquitylated uS3 and uS5 (Figure 5A) (Meyer et al., 2020). These modifications were not further induced upon HTN treatment, suggesting that loss of USP10 results in maximal uS3 and uS5 ubiquitylation which cannot be further augmented by the stressors used here (Figure 5A). Similarly, treatment with ISR agonists or high dose elongation inhibitors did not further elevate uS3 and uS5 ubiquitylation in USP10-KO cells (Fig. 5B). These observations suggest that preinitiation collisions are pervasive in the absence of stress, and that excess levels of USP10 relative to RNF10 maintain low levels of ribosomal ubiquitylation at basal conditions. In agreement with this, exogenous USP10 overexpression resulted in a loss of observable ribosomal ubiquitylation that was largely dependent upon the deubiquitylating activity of USP10 (Fig. 5A). Surprisingly, in USP10-KO cells, stress-induced uS3 and uS5 ubiquitylation decreased at later timepoints (Fig 5B). Because these cells lack the principle uS3 and uS5 deubiquitylating enzyme, the observed loss of ribosomal ubiquitylation was puzzling.

**Figure 5.**
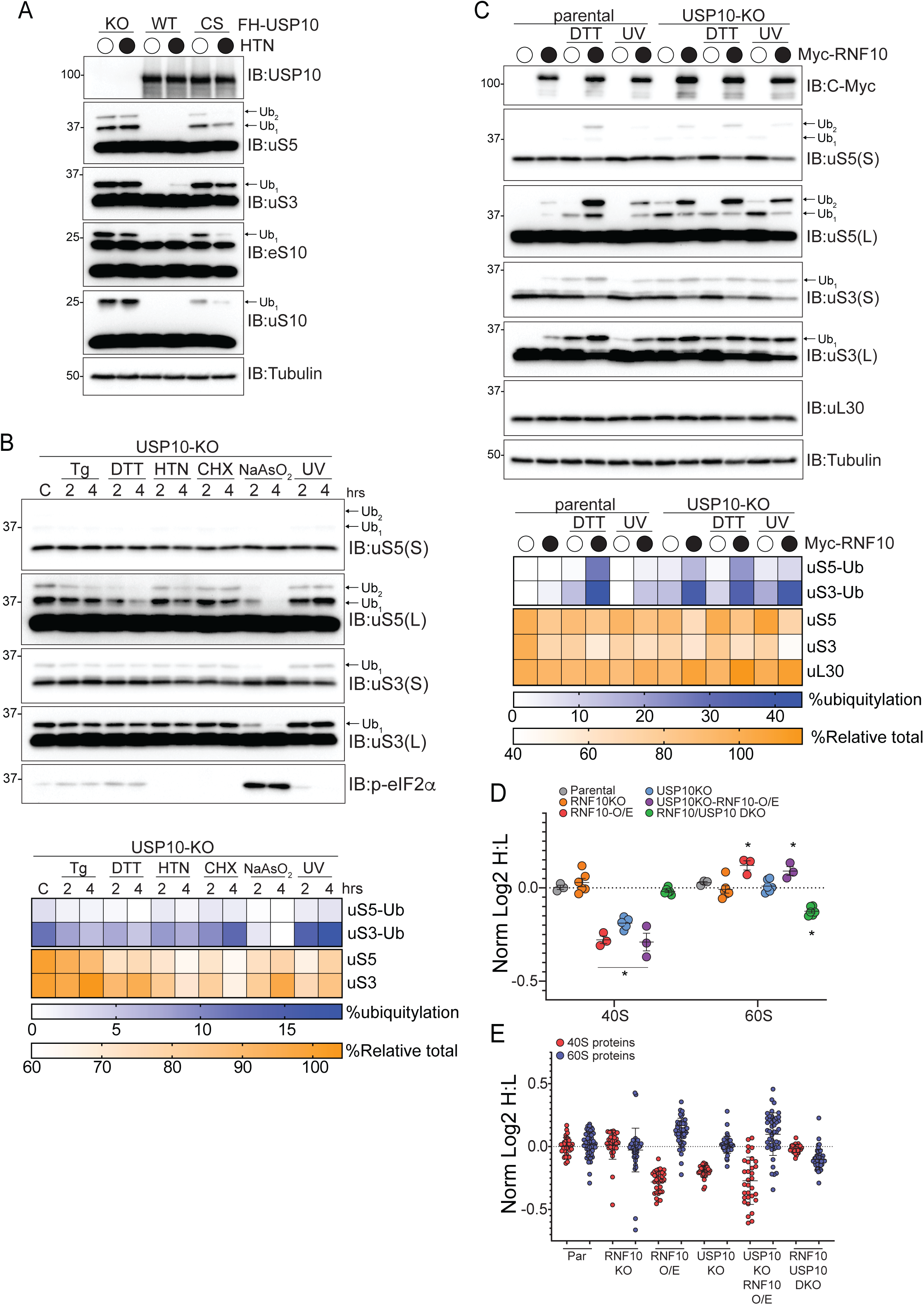
Persistent uS3 and uS5 ubiquitylation targets 40S ribosomal proteins for degradation, See also Figure S4. (A) 293T USP10-knockout (KO) cells constitutively expressing wild type (WT) or inactive mutant (CS) USP10 were treated with HTN (2ug/ml) for 2 hours and cell lysates were analyzed by SDS-PAGE and immunoblotted with the indicated antibodies. For all blots the ubiquitin-modified ribosomal protein is indicated by the arrow. S and L denote short and long exposures, respectively. (B) (top) USP10-KO cells were treated as indicated and cell extracts were analyzed by SDS-PAGE and immunoblotted with the indicated antibodies. (bottom) Percent ubiquitylated uS3 and uS5, and percent total relative abundance quantified from immunoblots (C) (top) Parental 293T or USP10-KO cells expressing Myc-tagged wild type RNF10 were either untreated, treated with DTT (5mM), or exposed to UV (0.02J/cm2). Cell lysates were analyzed by SDS-PAGE and immunoblotted with the indicated antibodies. (bottom) Quantitative representation of uS3 and uS5 percent ubiquitylation, and percent relative total abundance for uS3, uS5 and uL30. (D) The median normalized log2 SILAC ratio (H:L) for all quantified 40S and 60S ribosomal proteins comparing parental cells (light label) to cells of the indicated genotype (heavy label) with or without RNF10 overexpression (O/E). Each point represents a biological replicate, Bars denote mean value for replicate experiments with error bars displaying SEM. *=pvalue<0.05 by student’s t test compared to parental controls. (E) The median normalized log2 SILAC ratio (H:L) for individual 40S and 60S ribosomal proteins comparing parental cells (light label) to cells of the indicated genotype with or without RNF10 overexpression (O/E). Bars denote mean and error bars denote SD.

### iRQC acts post-translationally to reduce 40S abundance

We surmised that the observed loss in uS5 and uS3 ubiquitylation upon protein synthesis inhibition was due to protein degradation in the absence of new protein production. To examine this possibility, we overexpressed RNF10 in the presence and absence of USP10, followed by ISR activation (Figure 5C). RNF10 overexpression in parental cells resulted in robust uS3 and uS5 ubiquitylation, with a notable increase in the extent of di-ubiquitin modified uS5 (Figure 5C). Again, we observed a noticeable reduction in overall uS3 and uS5 protein levels in DTT-treated cells overexpressing RNF10 (Figure 5C). These observations were further enhanced when we overexpressed RNF10 in USP10 knockout cells, both at steady state and following stress induction (Figure 5C). In contrast, levels of the 60S subunit protein uL30 (RPL7) were unchanged upon RNF10 overexpression in either parental or USP10-KO cells. To examine if RNF10 expression suppressed ribosomal gene transcription, we measured uS3, eS6 (RSP6), or uL30 mRNA abundance in cells overexpressing RNF10. RNF10 overexpression did not decrease mRNA abundance, consistent with a post-transcriptional mechanism underlying the observed reduction in 40S protein levels (Figure S4B). Taken together, these results suggest that the abundance of 40S, but not 60S, ribosomal proteins is decreased upon conditions that result in constitutively high levels of uS3 and uS5 ubiquitylation.

To determine if the observed reduction in 40S protein levels occurred due to lower mRNA translation or elevated protein degradation, we used a metabolic pulse labeling approach: heavy SILAC-labeled cells were switched to the light label and ribosomal protein synthesis was followed over time by quantitative proteomics. Global protein synthesis rates were unaltered in USP10 KO cells whereas a small but significant increase occurred in the rate of 40S, but not 60S, protein synthesis (Figure S4C). RNF10 overexpression suppressed global protein synthesis rates, consistent with observations that RNF10 overexpression reduces cellular proliferation rates. Despite this decrease in overall protein synthesis, 40S protein synthesis rates were increased in cells overexpressing RNF10 relative to 60S or total protein synthesis rates (Figure S4C). These results indicate that the observed selective reduction in 40S compared to 60S protein levels when uS3 and uS5 ubiquitylation is enhanced is not due to a decrease in 40S protein synthesis. In fact, cells with RNF10 overexpression appear to activate mechanisms to offset reduced 40S protein abundance by increasing 40S protein synthesis rates.

### iRQC pathway activation results in 40S protein degradation

To examine if enhanced uS3 and uS5 ubiquitylation results in the specific loss of uS3 and uS5 protein, or more broadly impacts overall 40S ribosomal protein abundance, we used SILAC-based quantitative proteomics to compare ribosome protein levels between parental cells and RNF10-KO, USP10-KO, USP10/RNF10 double knockout cells, or cells overexpressing RNF10. RNF10-KO cells had comparable 40S and 60S protein levels to parental cells, whereas RNF10 overexpression resulted in a ∼17% reduction in 40S ribosomal protein abundance while modestly increasing 60S protein levels (Figure 5D; Table S2). Consistent with a previous report, cells lacking USP10 have reduced 40S protein levels (Meyer et al., 2020). Furthermore, 60S protein levels were unchanged relative to parental cells in USP10-KO cells. RNF10 overexpression in USP10-KO cells further reduced 40S protein levels while slightly increasing 60S protein abundance (Figure 5D). The observed decrease in 40S protein abundance in UPS10-KO cells was reversed in RNF10/USP10 double KO cells indicating that RNF10-dependent uS5 and uS3 ubiquitylation promotes 40S protein loss in USP10-KO cells. The abundance of the entire 40S subunit, rather than individual proteins, is reduced upon RNF10 overexpression or USP10 depletion suggesting that overall 40S protein stability is reduced by uS3 and uS5 ubiquitylation (Figure 5E; Table S2). These results are consistent with a model where preinitiation complex collisions are ubiquitylated by RNF10, and those ubiquitylated 40S ribosomal subunits that escape USP10-dependent demodification are targeted for degradation.

### 40S protein degradation is autophagy independent

Overall, our data indicate that preinitiation collisions result in 40S degradation. An autophagic mechanism seemed most plausible given that previous studies in *S. cerevisiae* have demonstrated that starvation conditions that inhibit mTOR signaling and stimulate autophagic flux result in enhanced ribosomal turnover by the autophagy pathway (Kraft et al., 2008). While mTOR-dependent degradation of ribosomes via autophagy does not appear to play a large role in regulating ribosomal abundance in mammalian cells (An and Harper, 2018; An et al., 2020), we directly evaluated if uS3 or uS5 ubiquitylation resulted in autophagy-dependent degradation of 40S ribosomal subunits. We first examined uS3 and uS5 protein levels upon RNF10 overexpression in parental or autophagy-deficient cells that are devoid of the critical ULK1 complex member, RB1CC1 (FIP200) (An et al., 2019). USP10 depletion or RNF10 overexpression alone or in combination resulted in the expected increase in uS3 and uS5 ubiquitylation and loss in 40S protein abundance in both parental and RB1CC1 knockout cells (Figure 6A). These results establish that autophagy-deficient cells maintain the ability to degrade RNF10 targeted 40S proteins. To confirm that RNF10-mediated ribosome ubiquitylation does not target 40S proteins for autophagy-dependent degradation, we utilized cell lines in which the genomic loci of uS3 or eL28 (RPL28) were tagged with the pH-sensitive fluorophore, Keima. These cell lines have been previously used to characterize ribosomal protein flux through the autophagy pathway (An and Harper, 2018). Consistent with previous reports, inactivation of mTOR signaling enhanced both 40S and 60S flux to lysosomes as indicated by an increase in the red to green Keima fluorescence (Figure 6B) (An and Harper, 2018). However, RNF10 overexpression did not result in an increase in either 40S or 60S ribosomal flux to the lysosome (Figure 6B), further suggesting that ubiquitylated 40S subunits are not detectably targeted for degradation by the autophagy pathway.

**Figure 6.**
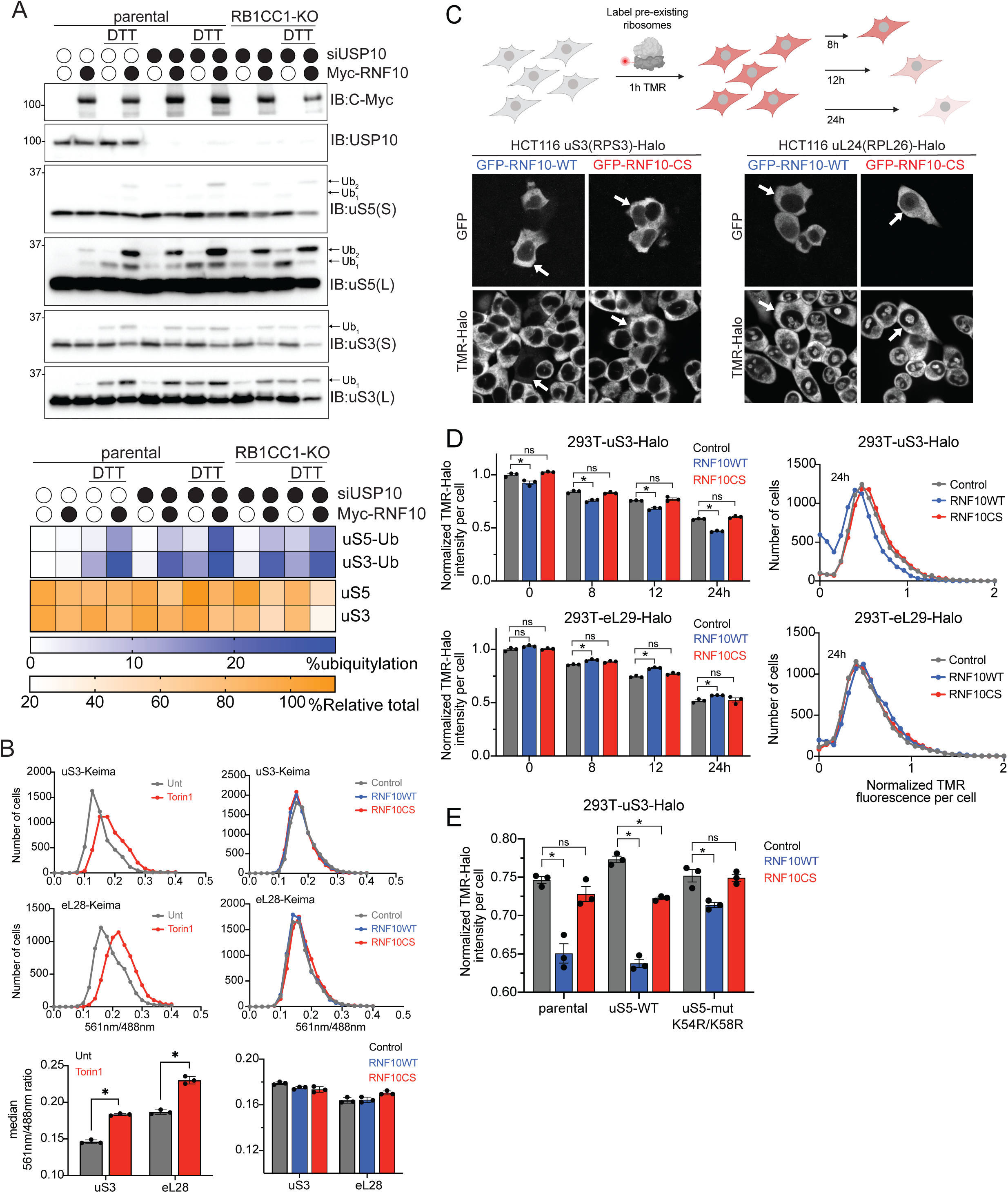
Enhanced ubiquitylation results in turnover of 40S ribosomal proteins in an autophagy-independent manner, See also Figure S4. (A) (top) Cell extracts from parental 293T or RB1CC1-KO cells transfected with either a control siRNA oligo or siRNA oligo targeting USP10, followed by transfection with Myc-tagged wild type RNF10 treated as indicated were analyzed by SDS-PAGE and immunoblotted with the indicated antibodies. (bottom) Quantitative representation of percent relative total abundance and uS3 and uS5 percent ubiquitylation. (B) HEK293 uS3 or eL28 (RPL28) Keima-tagged cells were either untreated (grey line) or Torin1 (150nM) treated (red line) for 24 hours. Cells expressing either a control plasmid (grey line), RNF10 wild type (blue line) or the catalytic mutant (red line) 48 hours post transfection were collected and analyzed via FACS. Frequency distributions of the red (561nm) to green (488nm) ratio are plotted. Bar graphs denote median red:green ratio from triplicate experiments. N=3, error bars denote SD of triplicate experiments. *=pvalue<0.0002 by unpaired student’s t test. (C) Schematic of Ribo-Halo fluorescent pulse-chase assay (top). Microscopy images of HCT116 uS3 or uL24-Halo tagged cells expressing GFP-tagged wild type (WT) or inactive mutant (CS) RNF10. Ribosomes were labeled with TMR ligand for 2 hours prior to imaging. Arrows indicate the same cells across panels (bottom). (D) The normalized (to control at 0h washout) TMR-fluorescence intensity for uS3 or eL29-Halo tagged cells expressing a control plasmid (grey bars), Myc-RNF10-WT (blue bars), or CS mutant (red bars) expression plasmid is depicted at the indicated time points post TMR washout (left). *=pvalue<0.05, ns = non-significant by multiple unpaired t tests compared to control protein. Frequency distribution of the normalized TMR signal at 24h is plotted (right). (E) TMR fluorescence intensities 12h post washout from parental 293T uS3-Halo tagged cells alone or with stable expression of wild type (WT) or K54R/K58R mutant (Mut) uS5 and transfected with a control plasmid (grey bars), GFP-RNF10-WT (blue bars), or GFP-RNF10-CS mutant (red bars) expression plasmids are depicted. The normalized (to control at 0h washout) TMR intensities are depicted. *=pvalue<0.05, ns = non-significant by multiple unpaired t tests compared to control protein.

### RNF10 mediated uS5 ubiquitylation accelerates 40S protein turnover

Ribosomal proteins are inherently stable and immunoblotting techniques can be insufficient to directly evaluate ribosome protein dynamics. In order to directly and quantitatively examine 40S and 60S protein degradation, we utilized previously characterized cell lines in which the genomic uS3, uL24 (RPL26), or eL29 (RPL29) loci were tagged with Halo (hereafter called Ribo-Halo) (An et al., 2020). These Ribo-Halo cell lines enable evaluation of ribosomal protein degradation kinetics through fluorescent pulse-chase experiments using fluorescently labeled Halo ligands (Figure 6C). Ribo-Halo cells overexpressing a control protein (LRRC49), wild type RNF10, or inactive RNF10 were red-labeled with a tetramethylrhodamine (TMR) Halo ligand for 1 hour to mark the existing pool of uS3, uL24, or eL29. Following TMR labeling, excess label was washed out and the abundance of the TMR-labeled ribosomal pool was monitored over time by microscopy. Three days post-transfection, cells expressing wild type GFP-RNF10 but not inactive GFP-RNF10CS displayed a marked decrease in cellular uS3-Halo abundance while having no impact on uL24-Halo protein levels (Figure 6C). To directly evaluate uS3 protein turnover, we performed pulse-chase experiments upon RNF10 expression and quantified single-cell Ribo-Halo abundance by flow cytometry. Ribo-Halo abundance was initially measured 36 hours post transfection, and ribosome decay was observed following TMR washout for 24 hours. These experiments revealed an increased uS3 turnover rate in cells expressing wild type RNF10 that was not observed in cells expressing a control protein or inactive RNF10 (Figure 6D). Consistent with our proteomics results, RNF10 overexpression did not increase turnover of the 60S subunit protein eL29 (Figure 6D). These results indicate that constitutive uS3 and uS5 ubiquitylation enhances 40S, but not 60S, protein degradation.

Our previous studies delineated a hierarchical relationship among uS3 and uS5 ubiquitylation events such that eliminating uS3 ubiquitylation renders uS5 incompetent for ubiquitylation, whereas eliminating uS5 ubiquitylation did not prevent uS3 ubiquitylation (Garshott et al., 2020). Based on these results, we engineered uS3-Halo cell lines to express either wild type or a ubiquitylation deficient mutant version (K54R;K58R) of uS5. Consistent with previous results, stable expression of exogenous uS5 comprised 80% of total uS5 levels (Figure S4D). While uS5 ubiquitylation in cells expressing exogenous wild type uS5 remained intact following DTT treatment, uS5 ubiquitylation was absent upon expression of mutant uS5 (Figure S4D). We then performed Ribo-Halo pulse-chase experiments in the uS3-Halo cells containing wild type or ubiquitylation mutant uS5. Consistent with previous results, wild type RNF10 overexpression resulted in an 18% decrease in uS3-Halo levels 12 hours post TMR washout in cells expressing wild type uS5, compared to cells expressing a control protein, (Figure 6E). However, RNF10 overexpression failed to accelerate uS3 degradation in cells expressing the ubiquitylation deficient version of uS5 (Figure 6E). These experiments causally link RNF10-dependent enhanced 40S protein degradation to the observed increase in ribosome ubiquitylation, and demonstrate that uS5 ubiquitylation is required for 40S turnover. Taken together, our results demonstrate that RNF10-dependent uS3 and uS5 ubiquitylation acts on preinitiation collisions to selectively target 40S subunit ribosomal proteins for degradation in an autophagy-independent manner.

## Discussion

Regulating ribosome traffic on mRNAs requires dynamic and coordinated control of translation initiation, elongation, and termination rates. Rapid rates of preinitiation complex loading and scanning relative to translation start would generate queues of potentially collided 43S preinitiation complexes within 5’UTRs. Evidence for such queueing has been demonstrated using in vitro translation systems and translation complex profile sequencing (TCP-seq) in yeast and human cells (Bohlen et al., 2020; Shirokikh et al., 2019; Sogorin et al., 2012; Wagner et al., 2020). Further, generating queues of preinitiation complexes using cycloheximide or insertion of an upstream open reading frame (uORF) resulted in alternative start codon utilization and translation recoding (Ivanov et al., 2018; Kearse et al., 2019). Consistent with these observations, we now demonstrate that preinitiation ribosome collisions occur during both stress conditions that slow scanning preinitiation complex progression, and -surprisingly - unperturbed cellular growth conditions

Seminal studies have long established that protein biogenesis activity is regulated by the presence of either structured elements or upstream uORFs within the 5’UTR of mRNAs (Ding et al., 2012; Gu et al., 2010; Hershey et al., 2019; Hinnebusch et al., 2016; Kozak, 1986a, b). Shorter, unstructured 5’UTRs enable more efficient translation, and secondary structures like G-quadruplexes and stable hairpins have been shown to robustly repress translation by inhibiting preinitiation complex scanning (Arora et al., 2008; Beaudoin and Perreault, 2010; Davuluri et al., 2000; Endoh and Sugimoto, 2016; Halder et al., 2009; Kozak, 1991; Sample et al., 2019). Further, multiple RNA helicases operate during translation initiation to assist in scanning through long and structured 5’UTRs (Gulay et al., 2020; Parsyan et al., 2011; Popa et al., 2016; Rubio et al., 2014; Sen et al., 2019; Sen et al., 2015; Soto-Rifo et al., 2012; Svitkin et al., 2001; Waldron et al., 2019; Wolfe et al., 2014). Despite the well-characterized repression of protein biogenesis associated with impeding progression of scanning ribosomes, the molecular rationale for this repression remained uncharacterized. Our data suggests that mRNAs containing structured 5’UTRs may increase the frequency of preinitiation complex collisions that terminate initiation and catalyze 40S degradation.

Collectively, our results demonstrate that ribosome collisions, both during the elongation and initiation phases of translation, nucleate site-specific 40S regulatory ribosomal ubiquitylation to modulate translation activity. The similar structural location and hierarchical relationship of 40S ribosomal proteins that are ubiquitylated during the elongation (uS10, eS10) and initiation (uS5, uS3) RQC pathway activation implies that the function of ribosome ubiquitylation may also be shared. However, the precise role ribosomal ubiquitylation plays during eRQC remains mysterious. Contrary to what is reported here for iRQC ubiquitylation events, elevating the extent of uS10 and eS10 ubiquitylation by overexpressing ZNF598 or removing deubiquitylating enzymes that antagonize ZNF598 does not result in detectable 40S degradation (Garshott et al., 2020).

Because RNF10-catalyzed 40S degradation is autophagy-independent, 40S degradation, in a presumably proteasome-dependent manner, would require 40S disassembly prior to degradation. Thus, iRQC-dependent ribosomal degradation appears distinct from the proteasomal degradation of unassembled ribosomal proteins mediated by either Huwe1 or Ube2O (Nguyen et al., 2017; Sung et al., 2016; Yanagitani et al., 2017). However, it is possible that the factors and mechanisms by which the 40S subunit is disassembled and degraded during iRQC are shared by those that target orphan ribosomal proteins for destruction.

Interestingly, in cells lacking USP10, uS5 and uS3 ubiquitylation reaches 20% of total uS5 and uS3 protein. These levels approach and surpass what has been observed for histone ubiquitylation, the most abundantly, and originally identified, ubiquitylated protein in the cell. The large extent of ribosome ubiquitylation in USP10-KO cells also suggests that preinitiation collisions occur at a high frequency in unstressed, albeit rapidly dividing, cells and that USP10 rapidly reverses ubiquitylation of these collied preinitiation complexes. The fact that uS3 and uS5 ubiquitylation is low in cells with USP10 argue that translation activity has evolved to allow for rapid translation initiation rates and subsequent collisions by maintaining an excess of USP10 relative to RNF10 (Nusinow et al., 2020) (Figure 7). Further, controlling the relative USP10:RNF10 ratio would set the threshold for collision tolerance at steady state while enabling stress-sensitive collision responses. These observations also suggest that, with sufficient USP10 activity, collided preinitiation complexes can eventually transition into elongating ribosomes (Figure 7).

**Figure 7.**
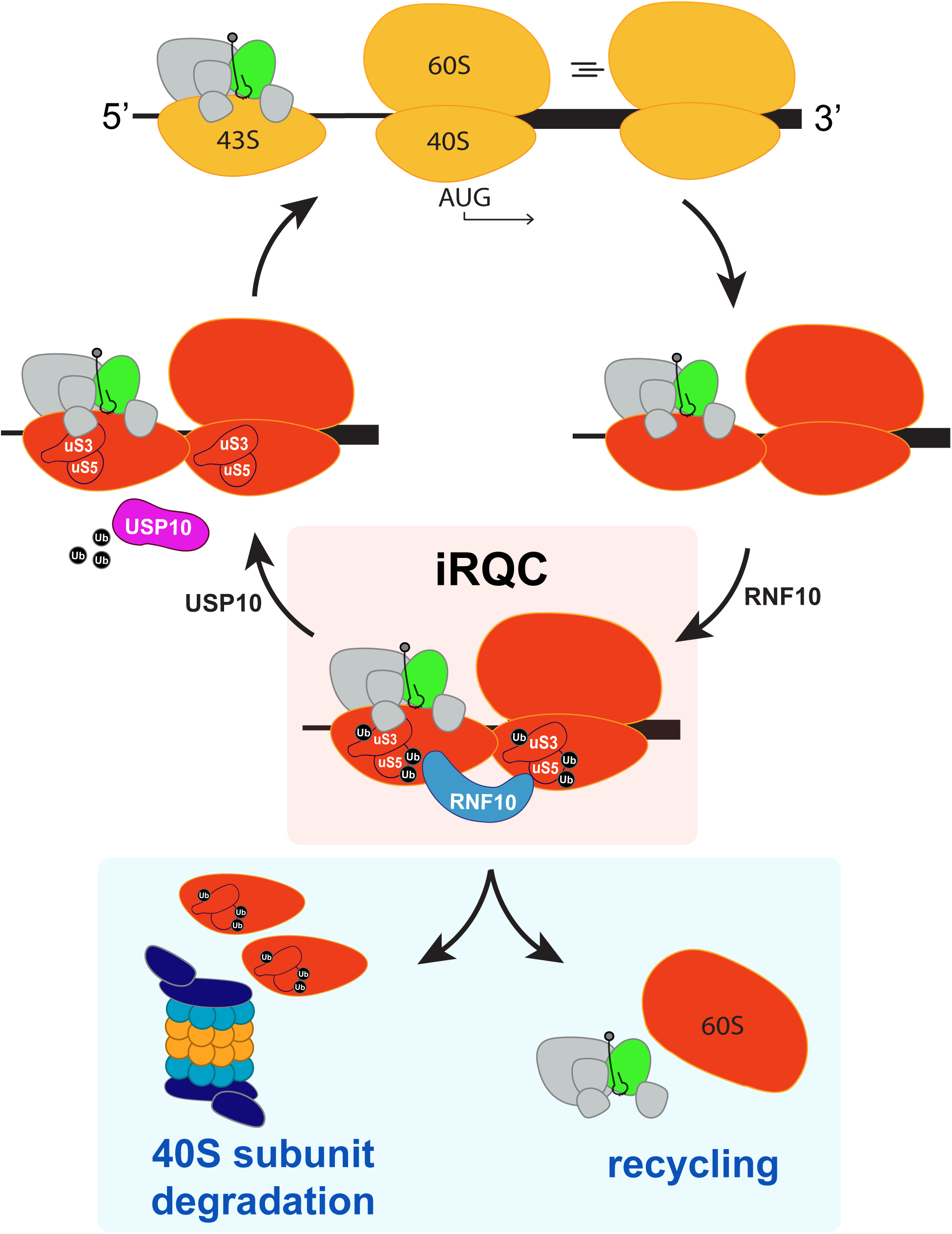
Model of iRQC activation following preinitiation complex collisions. Under normal homeostatic conditions, cap-dependent translation proceeds with 43S scanning of the 5’ untranslated region, followed by start codon recognition, and 60S subunit joining to form an elongation competent 80S ribosome (indicated by yellow ribosomes). Collisions between 43S and 80S ribosomes (red ribosomal subunits) activate the iRQC pathway in which RNF10 is recruited to collisions involving preinitiation complexes and ubiquitylates uS5 and uS3 on specific lysine residues. Persistent ribosome ubiquitylation triggers 40S subunit destruction and recycling of initiation factors and the 60S subunit. Transient collisions can undergo USP10-dependent ribosome deubiquitylation to allow for translation initiation progression

Similar to eRQC, the iRQC pathway appears to be conserved in single-celled eukaryotes as USP10 and RNF10 orthologs have been shown to regulate uS3 ubiquitylation in *S. cerevisiae* (Jung et al., 2017; Sugiyama et al., 2019). Not only are the enzymes conserved, but so too are the mechanistic requirements: yeast with inactivating mutations in the peptidyl transferase center of ribosomes that allow for scanning, but block elongation, trigger ribosomal RNA decay in a manner dependent upon uS3 ubiquitylation (Sugiyama et al., 2019). Interestingly, and completely opposite to what we observed during mammalian iRQC, the USP10 homolog in *S. cerevisiae*, Ubp3, has also been implicated in regulating 60S, but not 40S, ribosome degradation in an autophagy-dependent manner upon starvation (Kraft et al., 2008). Future studies are needed to disentangle starvation-induced ribophagy from iRQC-mediated 40S degradation as they appear to utilize overlapping components.

We propose that conserved ribosomal ubiquitylation acts as a cellular rheostat to dynamically regulate translation dynamics during conditions that enhance collision frequencies. Accumulating evidence suggests that elongation collisions not only trigger ribosomal subunit recycling, but also throttle translation initiation rates (Meydan and Guydosh, 2020; Vind et al., 2020). However, previous studies did not observe whether ribosome collisions occurred prior to translation initiation and what their impact would be on translation activity. Our data establish that preinitiation collisions trigger 40S subunit degradation which would serve to globally reset translation initiation rates. As cellular proliferation rates change, for example during cellular differentiation, translation capacity and ribosome abundance may also be altered to match metabolic needs. An increase in preinitiation collision frequencies may signal ribosome overabundance which could be rectified by iRQC-mediated degradation. Our demonstration that ISR-stimulating conditions also induce preinitiation collisions suggests that chronic stress signaling may also reset translation capacity. These findings describe a previously uncharacterized, and likely conserved, distinct ribosome-associated quality control pathway that can be utilized to regulate 40S ribosomal levels.

## Acknowledgments

We thank Jody Puglisi and Alex Johnson (Stanford University) for technical advice and for providing purified ribosomal submits. We also acknowledge Julie Monda, Xuezhen Ge, and Steven Wasserman (UCSD) for editing assistance during manuscript preparation. This research was supported by funding from the National Institutes for Health (DP2GM119132 (E.J.B.), R01GM136994 (E.J.B.), R01AG011085 (J.W.H.), R01NS083524 (J.W.H.), T32GM007240 (D.M.G.), and the National Science Foundation (DGE-1650112 (D.M.G.)).

## Author contributions

Conceptualization, D.M.G. and E.J.B.; Methodology, D.M.G., H.A., and E.J.B.; Investigation, D.M.G., H.A., E.S., M.L., A.V. and E.J.B.; Visualization, D.M.G., H.A. and E.J.B.; Funding Acquisition, E.J.B and J.W.H.; Supervision, J.W.H., and E.J.B.; Writing-original draft, D.M.G. and E.J.B.; Writing-Review & Editing, D.M.G., H.A., E.S., M.L., A.V., J.W.H. and E.J.B.

---

**Figure S1.**
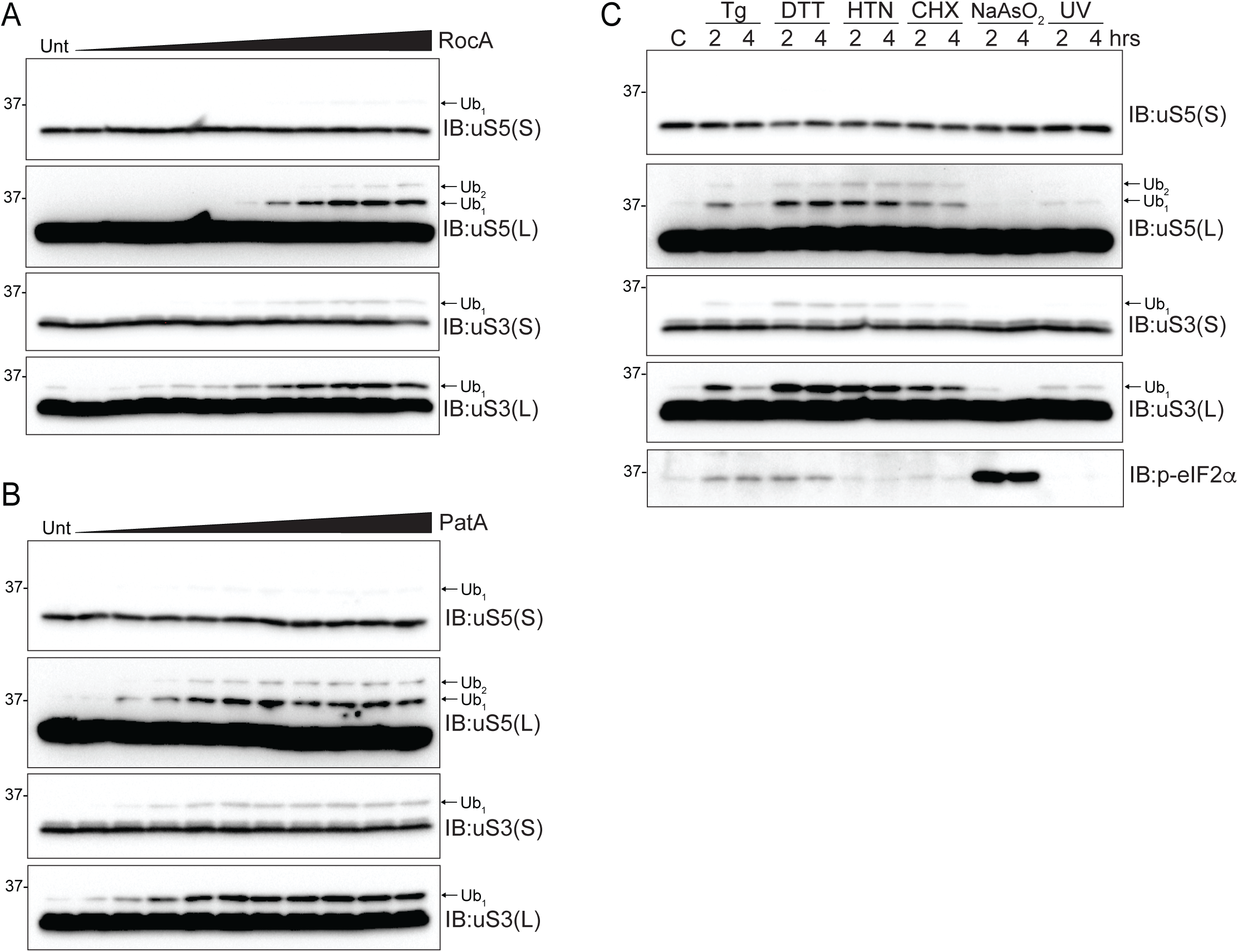
Translation initiation inhibition triggers preinitiation complex collisions and ribosome ubiquitylation, Related to Figures 1 and 2. (A,B) 293T cells were treated with increasing concentrations of either RocA (0.031-3.2uM) (A) or PatA (0.31-100nM) (B) for two hours. Whole-cell lysates were analyzed by SDS-PAGE and immunoblotted with the indicated antibodies. For all blots, the ubiquitin-modified ribosomal protein is indicated by the arrow. S and L denote short and long exposures, respectively. (C) Cell extracts from cells treated with Tg (1uM), DTT (5mM), HTN (2ug/ml), CHX (100ug/ml), NaAsO2 (500uM) or UV (0.02J/cm2) (washout) for two or four hours were analyzed by SDS-PAGE and immunoblotted with the indicated antibodies.

**Figure S2.**
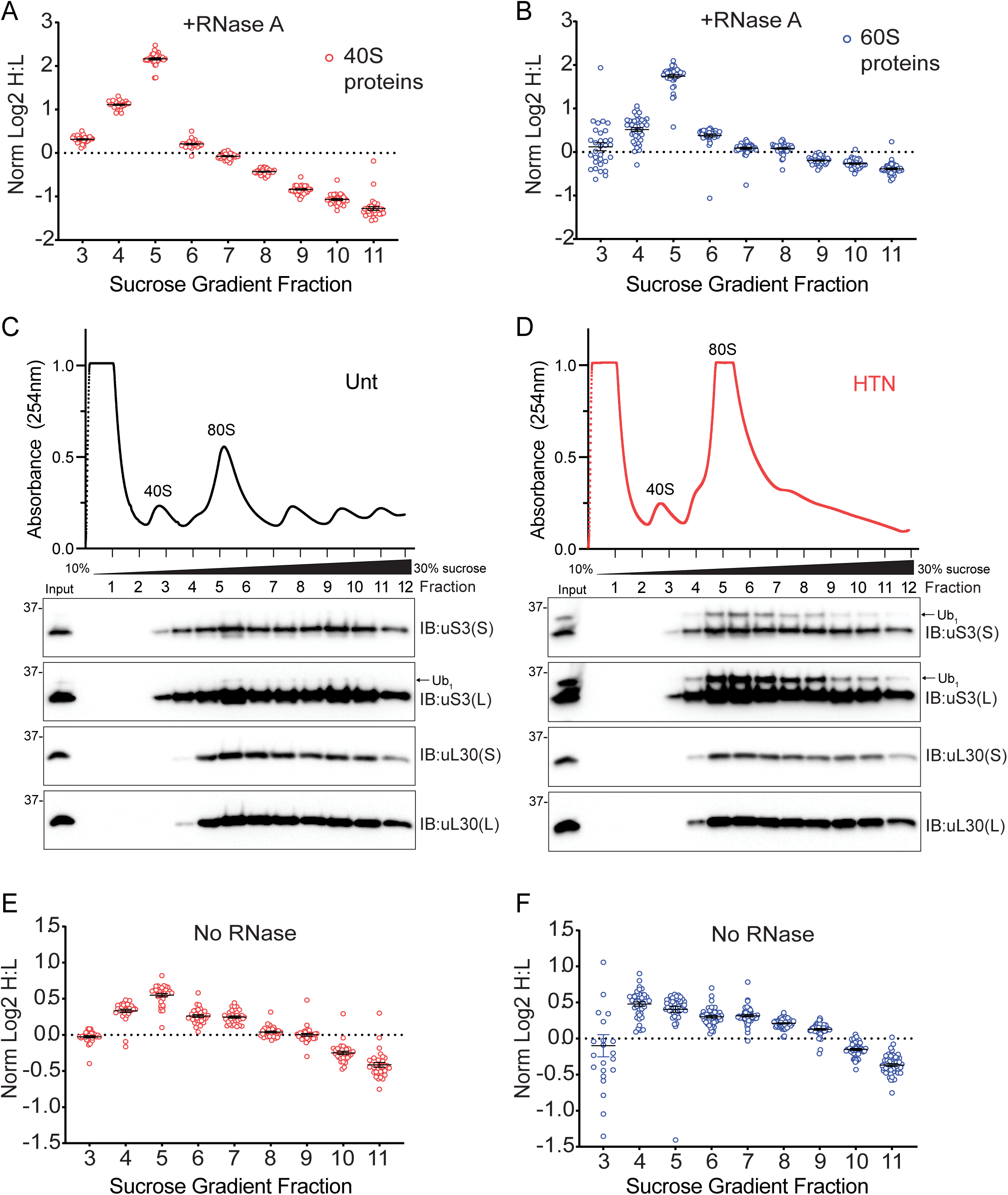
Ribosomal subunit imbalance is present in gradient fractions containing ubiquitylated ribosomes, Related to Figure 3. (A-B) The normalized log2 SILAC ratio (H-HTN:L-untreated) of all quantified 40S (A, red circles) and 60S (B, blue circles) proteins within indicated sucrose gradient fractions from lysates treated with RNaseA prior to density gradient centrifugation. Each data point is an individual ribosomal protein, and the black bar denotes the median value. (C-D) Cell extracts from untreated (C) or HTN (2ug/ml) treated (D) 293T cells were fractionated on 10-30% sucrose gradients. The 254nm absorbance trace is depicted. Fractions were analyzed by SDS-PAGE and immunoblotted with the indicated antibodies. The ubiquitin-modified ribosomal protein is indicated by the arrow. S and L denote short and long exposures respectively. (E-F) The normalized log2 SILAC ratio (H-HTN:L-untreated) of all quantified 40S (E, red circles) and 60S (F, blue circles) proteins within indicated sucrose gradient fractions from untreated lysates prior to density gradient centrifugation. Each data point is an individual ribosomal protein, and the black bar denotes the median value.

**Figure S3.**
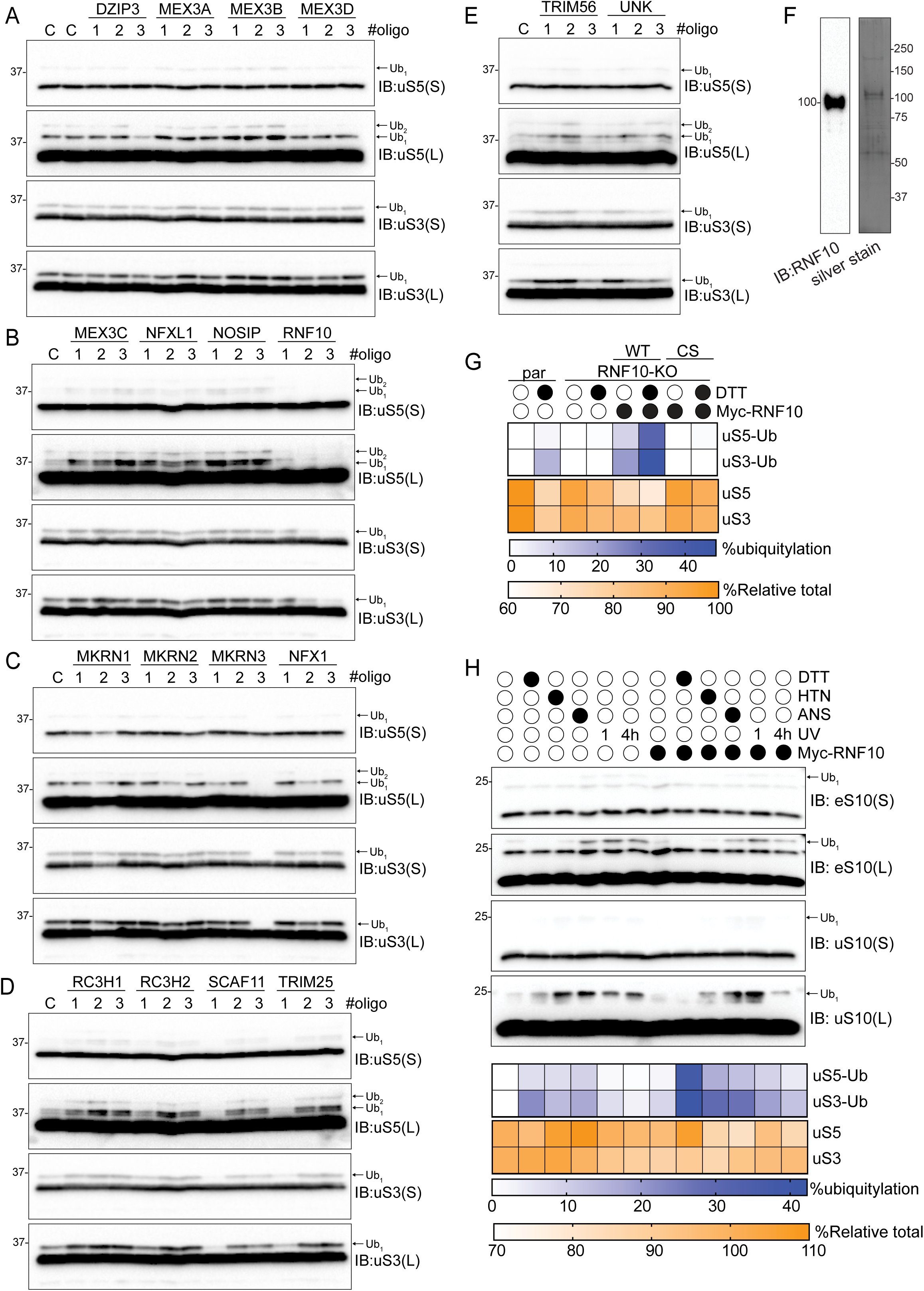
Screening RNA-associated ubiquitin ligases identifies RNF10 as the uS3/uS5 ubiquitin ligase, Related to Figure 4. (A-E) 293T cells were transfected with either a control siRNA oligo or three separate siRNA oligos targeting the indicated E3 ligase, followed by treatment with DTT (5mM) for two hours. Cell lysates were analyzed by SDS-PAGE and immunoblotted with the indicated antibodies. For all blots, the ubiquitin-modified ribosomal protein is indicated by the arrow. S and L denote short and long exposures, respectively. (F) RNF10 immunoblot and silver stain of affinity purified wild type RNF10 expressed in 293T cells. (G) Quantitative representation of percent ubiquitylated uS3 and uS5, and percent relative total abundance from immunoblots in 4E. (H) (top) 293T cells with and without Myc-tagged wild type RNF10 expression were drug treated as indicated. UV indicates that cells were exposed to UV (0.02J/cm2) and were allowed to recover for 1 or 4 hours. Cell extracts were analyzed by SDS-PAGE and immunoblotted with the indicated antibodies. (bottom) Quantitative representation of uS3 and uS5 percent ubiquitylation and percental relative total abundance from immunoblots in 4F.

**Figure S4.**
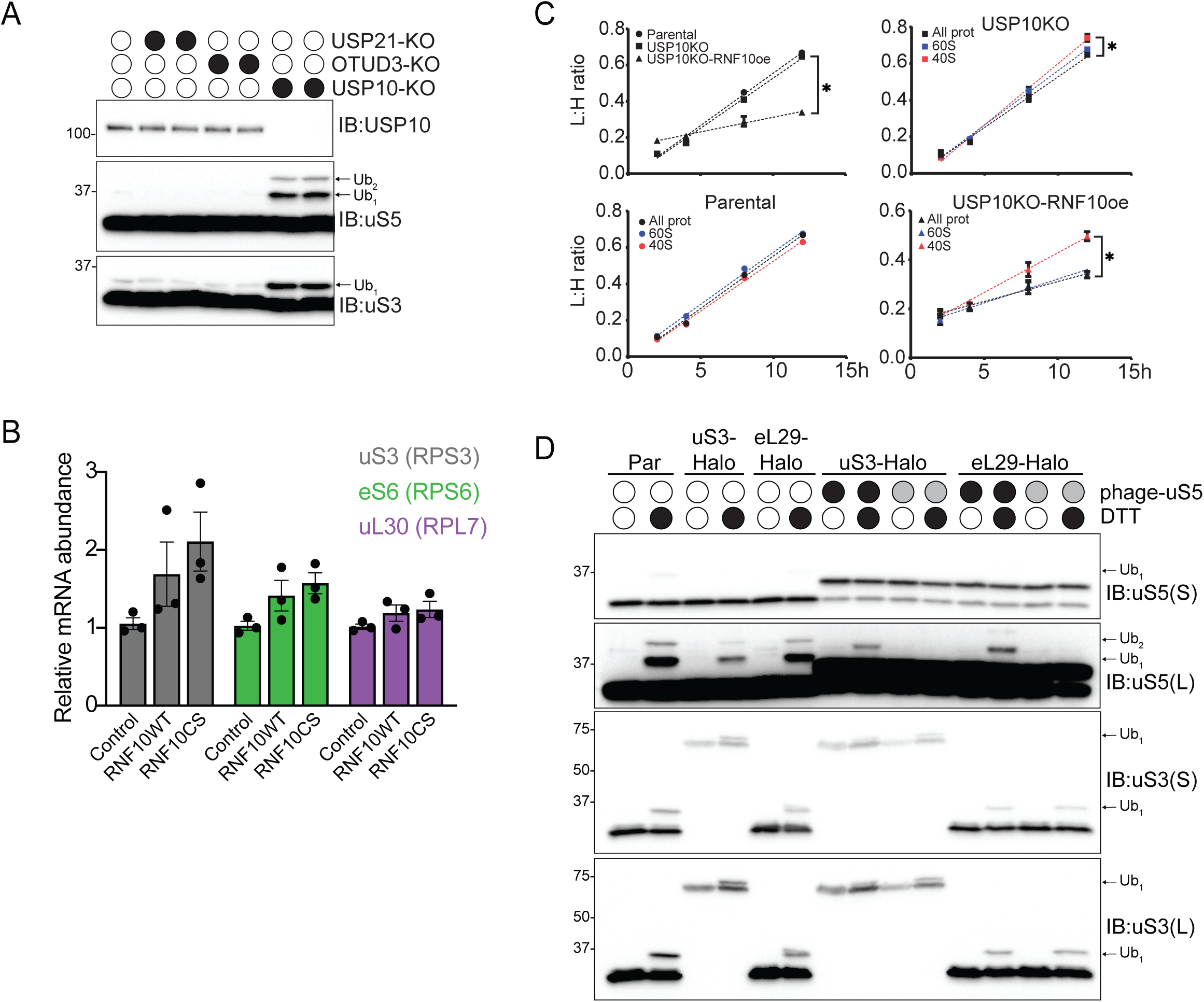
uS3 and uS5 ubiquitylation trigger 40S subunit turnover, Related to Figures 5 and 6. (A) Cell extracts from parental 293T cells or USP21-KO, OTUD3-KO or USP10-KO cells were analyzed by SDS-PAGE and immunoblotted with the indicated antibodies. For all blots, the ubiquitin-modified ribosomal protein is indicated by the arrow. S and L denote short and long exposures, respectively. (B) Relative mRNA abundance measured by qPCR for uS3 (RPS3), eS6 (RPS6), and uL30 (RPL7) in parental cells, or parental cells expressing wild type (WT) or inactive mutant (CS) RNF10. N=3, error bars denote SEM. (C) Total protein (black line), 40S (red line), or 60S (blue line) ribosomal protein synthesis rates from parental 293T, USP10-KO, and USP10-KO cells overexpressing wild type RNF10 were determined using SILAC label swap from cells collected at the indicated time point post label swap. Light to heavy ratios were determined by mass spectrometry. N=3, error bars denote SEM of triplicate experiments. *=pvalue<0.05 using unpaired, two-tailed Student’s t-test comparing slope of best-fit line for replicate experiments. (D) 293T-uS3-Halo cells alone or with constitutive wild type (black circles) or K54R/K58R mutant (grey circles) uS5 expression were either untreated or treated with DTT (5mM) for 2 hours. Cell lysates were analyzed by SDS-PAGE and immunoblotted with the indicated antibodies.

**Table S1. Proteomic analysis of sucrose gradient fractions with and without HTN, Related to** Figures 3 and S2. Log2 SILAC ratios (H-HTN:L-unt) or label free quantification (LFQ) ion intensities for all measured ribosomal proteins within individual sucrose gradient fractions from untreated or RNase treated cell lysates across three replicate experiments. Data represented in Figures 3D and 3E and S2A-S2F.

**Table S2. Proteomic analysis of ribosome abundance in cells with altered iRQC activity, Related to** Figure 5. Log2 SILAC ratios (H-indicated genotype:L-parental) for all measured ribosomal proteins from parental, RNF10-KO, parental with RNF10 overexpression (OE), USP10-KO, USP10-KO with RNF10 OE, or USP10/RNF10 double KO (DKO) cells across replicate experiments. Data represented in Figures 6D and 6E.

## STAR METHODS

## RESOURCE TABLE

| REAGENT or RESOURCE | SOURCE | IDENTIFIER |
| --- | --- | --- |
| Antibodies |  |  |
| Rabbit monoclonal anti-RPS2 (uS5) | Bethyl Laboratories | Cat# A303-794A;<br>RRID:AB_11218192 |
| Rabbit monoclonal anti-RPS3 (uS3) | Bethyl Laboratories | Cat# A303-840A;<br>RRID:AB_2615588 |
| Rabbit polyclonal anti-RPS10 (eS10) | ABclonal | Cat# A6056;<br>RRID:AB_2766730 |
| Rabbit monoclonal anti-RPS20 (uS10_ | Abcam | Cat# ab133776;<br>RRID:AB_2714148 |
| Rabbit monoclonal anti-p-eIF2α | Cell Signaling Technology | Cat# 3398S;<br>RRID:AB_2096481 |
| Rabbit polyclonal anti-RPL7 (uL30) | Bethyl Laboratories | Cat# A300-741A;<br>RRID:AB_2301241 |
| Rabbit polyclonal anti-USP10 | Abcam | Cat# ab72486;<br>RRID:AB_1271412 |
| Mouse monoclonal α-Tubulin | Cell Signaling Technology | Cat# 3873;<br>RRID:AB_1904178 |
| Mouse monoclonal c-Myc | Santa Cruz Biotechnology | Cat# sc-40 AC;<br>RRID:AB_2857941 |
| Mouse monoclonal anti-tubulin | Cell Signaling Technology | Cat# 3873S; RRID:AB_1904178 |
| Anti-Rabbit IgG (H+L), HRP Conjugate antibody | Promega | Cat# W4011; RRID:AB_430833 |
| Anti-Mouse IgG (H+L), HRP Conjugate antibody | Promega | Cat# W4021; RRID:AB_430834 |
| Lipofectamine RNAiMax | Thermo-Fisher | Cat# 13778030 |
| Lipofectamine 2000 | Thermo-Fisher | Cat# 11668019 |
| Mirus TransIT 293 | Mirus Bio llc | Cat# MIR 2700 |
| Protease inhibitor cocktail tablet | Roche | Cat# 11836170001 |
| Immun-Blot® PVDF Membrane | BioRad | Cat# 1620177 |
| Albumin Standard | Thermo Scientific | Cat# 23209 |
| Clarity™ Western ECL Substrate | BioRad | Cat# 170-5061 |
| Chemicals |  |  |
| DL-Dithiothreitol | ACROS organics | Cat# 165680050 |
| Anisomycin | Fisher Scientific | Cat# 50995788 |
| Harringtonine | LKT labs | Cat# H0169 |
| Thapsigargin | Tocris Biosciences | Cat# 1138 |
| Cycloheximide | MP Biomedicals | Cat# 100183 |
| Sodium arsenite solution | Fluka Analytical | Cat# 35000-1L-R |
| Torin1 | Tocris Biosciences | Cat# 4247 |
| PatamineA | Gift from Jeremy Pelletier's lab |  |
| MG-132 | Enzo Life Sciences | Cat# BML-PI102 |
| Rocaglamide | MedChemExpress | Cat# HY19356 |
| Lactimidomycin | Millipore Sigma | Cat# 5062910001 |
| Doxycycline hydrochloride | Fisher Scientific | Cat# BP2653-5 |
| Ammonium persulfate ($\geq 98\%$) | Sigma | Cat# A3678 |
| Fetal Bovine Serum | VWR | Cat# 97068-085 |
| Trypsin | Sigma-Aldrich | Cat# T1426 |
| N-Ethylmaleimide | Sigma-Aldrich | Cat# E3876 |
| Puromycin | Mediatech<br>(Corning) | Cat# 61-385-RA |
| RNaseA | Thermo Scientific | Cat# EN0531 |
| SUPERase-In™ | Invitrogen | Cat# AM2696 |
| Turbo™ DNase | Invitrogen | Cat# AM2238 |
| Trizol | Life Technologies | Cat# 15596026 |
| Dimethyl sulfoxide | Corning | Cat# 25-950-CQC |
| β-mercaptoethanol | J.T. Baker | Cat# 4049-00 |
| Triton-X 100 | Sigma | Cat# T8787 |
| EDTA | Amersco | Cat# E177 |
| Trichloroacetic acid solution | Sigma | Cat# T0699 |
| Thermostable Inorganic<br>Pyrophosphatase | New England<br>Biolabs | Cat# M0296S |
| ATP solution | AMSBIO | Cat# 2121-100 |
| Creatine kinase | Biovision | Cat# 76289-160 |
| Creatine phosphate | Sigma Aldrich | Cat# 10621714001 |
| Critical Commercial Assays |  |  |
| BCA Protein Assay | Thermo Scientific<br>(Pierce) | Cat# 23225 |
| Bradford Assay; protein assay dye<br>reagent concentrate | BioRad | Cat# 500-0006 |
| HaloTag®TMR ligand | Promega | Cat# G8251 |
| Experimental Models: Cell Lines |  |  |
| 293T | ATCC | Cat# CRL-3216 |
| HCT116 | ATCC | Cat# CCL-247 |
| HEK293 | ATCC | Cat# CRL-1573 |
| 293Flp-In |  |  |
| 293T-USP10KO | This paper | N/A |
| 293T-USP21KO | Garshott et al. 2020 | N/A |
| 293T-OTUD3KO | Garshott et al. 2020 | N/A |
| HCT116-uS3-Halo | An et al. 2020 | N/A |
| HCT116-uL26-Halo | This paper | N/A |
| 293T-uS3-Halo | An et al. 2020 | N/A |
| 293T-eL29-Halo | An et al. 2020 | N/A |
| HEK293-uS3-Keima | An et al. 2018 | N/A |
| HEK293-eL28-Keima | An et al. 2018 | N/A |
| HEK293T-RB1CC1KO | An et al. 2020 | N/A |
| 293FlpIn-FRT-Flag-HA-RNF10wt | This paper | N/A |
| 293FlpIn-RNF10KO | This paper | N/A |
| 293FlpIn-RNF10/USP10-DKO | This paper | N/A |
| 293T-Flag-HA-USP10wt | This paper | N/A |
| 293T-Flag-HA-USP10CS | This paper | N/A |
| Recombinant DNA |  |  |
| pCMV-N-Myc-RNF10wt | This paper | N/A |
| pCMV-N-Myc-RNF10CS | This paper | N/A |
| pCMV-N-Myc-LRRC49 | This paper | N/A |
| pCMV-N-GFP-RNF10wt | This paper | N/A |
| pCMV-N-GFP-RNF10CS | This paper | N/A |
| pCMV-N-Myc-GFP | This paper | N/A |
| Sequence-Based Reagents |  |  |
| See Supplemental Experimental Procedures for siRNA reagents |  |  |
| Software and Algorithms |  |  |
| FlowJo (v10.6.2) | BD biosciences |  |
| Prism 9.0 | GraphPad software |  |

## CONTACT FOR REAGENT AND RESOURCE SHARING

Requests for resources or further information can be directed to the Lead Contact Eric J. Bennett.

## SUPPLEMENTAL EXPERIMENTAL PROCEDURES

### Plasmids

Using Gateway cloning (Invitrogen) all protein coding regions were cloned into Myc-of GFP-tagged CMV expression vectors. Mutations were introduced using QuickChange site-directed mutagenesis utilizing PCR-based approaches (primers 5’ to 3’: RNF10-C225S, CATGAAGTGCCATCTTCCCCAATATGCCTCTATC). Template DNA was digested by Dpn1 followed by transformation of the mutated plasmids into TOP10 E. coli cells. Plasmids were confirmed by sequencing and screened for expression by immunoblotting.

### Cell lines, transfections and siRNA

All HEK293, HEK293T, HCT116 and 293Flp-In cells were grown in DMEM (high glucose, pyruvate and L-Glutamine) containing 10% fetal bovine serum (FBS) and 1% penicillin/streptomycin and maintained in a 5% CO2 humidified incubator. Where indicated, before harvesting cells were treated with either 1uM Tg, 5mM DTT, 2ug/ml HTN, 100ug/ml CHX, 500uM NaAsO2, 150nM Torin1 or were exposed to 0.02J/cm2 UV radiation using a SpectorlinkerTM XL-1000 (Spectronics). Lentiviral transduction was used to generate stable cells lines expressing Flag-HA tagged USP10. Using Mirus TransIT 293 transfection reagent cells were transfected with five helper plasmids pHAGE-GAG-POL; pHAGE-VSVG; pHAGE-tat1b; pHAGE-rev and pHAGE-Flag-HA-USP10 (wild type or catalytic mutant), followed by the addition of fresh media after 24 hours. The supernatant was filtered using a 0.45 mm sterile syringe filter and mixed with 2ul of 6mg/ml polybrene. The viral mixture was then added to cells seeded at 50% confluency and infected for 24hours. Stable expression clones were selected with 1ug/ml Puromycin.

The Flp-In^TM^ system (Thermo Fisher) through single locus integration and hygromycin selection was used to generate stable doxycycline inducible cell lines expressing Flag-HA-tagged proteins. Flp-In 293 cells were transfected with Flp-In expression vectors for RNF10 using TransIT 293 transfection reagent (Mirus) according to manufacturer guidelines. Cells were seeded at 60% confluency, transfected for 24 hours followed by selection of stable expression clones with 100ug/mL Hygromycin. Treatment with 2ug/mL doxycycline for 16 hours prior to harvesting was used to induce protein expression.

All transient transfections were carried out using Lipofectamine 2000 (Thermo Fisher) and all siRNA knockdown transfections were performed using Lipofectamine RNAiMAX (Thermo Fisher) according to manufacturer instructions. A list of all RNAi oligonucleotides used in this study can be found in table below.

### Immunoblotting

For all immunoblot analysis, cell pellets were resuspended in urea denaturing lysis buffer (8M urea, 50mM Tris-Cl, pH 8.0, 75mM NaCl, 1mM NaV, 1mM NaF, 1mM β-glycerophosphate, 40mM NEM in the presence of EDTA-free protease inhibitor cocktail) and kept on ice during preparation. Cell lysates were sonicated for 10 s (output of 3W on a membrane dismembrator model 100 (Fisher Scientific) with a microtip probe then centrifuged for 10 min at 15,000rpm at 4°C. Lysate protein concentrations were measured by BCA Protein Assay (23225, Thermo Scientific Pierce). Laemmli sample buffer with β-mercaptoethanol was then added to cell lysates and heated at 95°C for 10 min. Samples were then cooled to room temperature and centrifuged briefly. Lysates were resolved on 12% Tris-glycine SDS -PAGE gels, followed by transfer to PVDF membranes (1620177, BioRad) using Bjerrum semi-dry transfer buffer (48mMTris Base, 39mM Glycine-free acid, 0.0375% SDS, 20% MeOH, pH 9.2) and a semi-dry transfer apparatus (Bio-Rad Turbo Transfer) for 30 min at 25V. Immunoblots were blocked with 5% blotting grade nonfat dry milk (APEX Bioresearch) in TBST for 1 hour. Primary antibodies were diluted in 5% BSA and rocked overnight. Immunoblots were developed using Clarity Western ECL Substrate (1705061, BioRad) and imaged on a Bio-Rad Chemi-Doc XRS+ system. All blots were processed using Imagelab (BioRad) software, with final images prepared in Adobe Illustrator. All plots were prepared using GraphPad Prism.

### Phos-Tag SDS-PAGE

For Phos-tag analysis, cell pellets were resuspended in 500ul of lysis buffer (8M urea, 50mM Tris-Cl, pH 8.0, 75mM NaCl, 1mM NaV, 1mM NaF, 1mM β-glycerophosphate in the presence of EDTA-free protease inhibitor cocktail). Lysates were sonicated for 10s (as described above) followed by centrifugation for 10 min at 15,000rpm at 4°C. 125ul of TCA was added to each sample, then incubated on ice for 2h at 4°C. Protein was collected by spinning tube in microcentrifuge at 15,000 rpm for 30min at 4°C. The TCA protein pellet was washed with 200ul cold acetone, followed by centrifugation at 15,000 rpm for 10min at 4°C. The acetone wash step was repeated two more times. Pellets were left to dry for 30min at room temperature to evaporate any remaining acetone, then resuspended in 50ul 8M urea/20mM DTT. Protein concentrations were measured by Bradford Assay (protein assay dye reagent concentrate, 500-0006, BioRad). Laemmli sample buffer with β-mercaptoethanol was then added to protein samples and heated at 95°C for 10 min. Samples were resolved on 12.5% SuperSepTM Phos-tagTM gels (198-17981, Fujifilm), followed by Zn2+ ion elimination. Gel was soaked in 1X transfer buffer (25mM Tris, 192mM Glycine, 10% v/v methanol) with 10mM EDTA for 20min with gentle agitation. This step was repeated three times with buffer exchanges, followed by 10min without EDTA. Wet-tank transfer to PVDF membranes using Towbin transfer buffer (25mM Tris, 192mM Glycine, 20% v/v methanol) was done overnight (16h) at 30V. Immunoblots were blocked, developed, and imaged as described above.

### Sucrose density gradient fractionation

Cell pellets were lysed in 500 ul of lysis buffer (20mM Tris-Cl, pH 8.0, 150mM NaCl, 15mM MgCl2, 1% Triton-X 100, 40U Turbo DNase I, 40mM NEM, 1mM DTT, EDTA-free protease inhibitor cocktail in DEPC treated water) followed by vigorous pipetting and incubated on ice for 15min. The cell lysates were centrifuged at 15,000 rpm for 10min at 4°C and the supernatant was transferred to a new microcentrifuge tube. Total RNA concentration of each lysate was determined using a nanodrop (Thermo Scientific). 500ug of total RNA was digested with 3.5ug/ml of RNaseA for 15min at 25°C on a thermomixer (Eppendorf) at 500rpm. The digestion was stopped with 166.5U of SUPERaseIn RNase Inhibitor. Samples were fractionated over a 10–30% sucrose gradient containing 150ug/ml cycloheximide (prepared on Gradient Master 108 (Biocomp): 1min 54s, 81.5 degrees, 16rpm). Samples were centrifuged at 41,000rpm for 2 hr at 4°C in an SW41i rotor. 1ml fractions were collected using a PGFip piston gradient fractionator (Biocomp). Protein fractions were precipitated overnight with 10% TCA at 4°C, followed by three ice-cold acetone washes. Pellets were dried in Vacufuge plus (Eppendorf) at room temperature for 5 min. Pellets were then resuspended in Laemmli sample buffer with β-mercaptoethanol, heated at 95°C for 10 min.

### SILAC LC-MS-MS analysis

Cells were grown in a media containing dialyzed FBS (FB03, Omega Scientific) and either light (K0) lysine and arginine (R0) or 13C615N2-labeled (K8) lysine and (R10) arginine (Cambridge Isotopes). Cells were harvested and mixed 1:1 by cell count and were processed for mass spectrometry as described previously (Markmiller et al., 2019). Briefly, cells were lysed using 8M urea lysis buffer with 40mM fresh NEM and lysates were quantified for protein content using the BCA assay. 20µg of total cell extract was diluted to a final urea concentration of 1M and then digested overnight with trypsin (V5111, Promega) at a 1:100 (enzyme:protein) ratio. The digests were reduced with 1mM DTT for 30 min and then alkylated with 10mM NEM in a dark for 30min. The digests were desalted using Stage-Tip method and analyzed by LC-MS/MS as described below. Mixed SILAC lysates were fractionated over sucrose gradients as described. Fractions were TCA precipitated, followed by resuspension in 50mM ammonium bicarbonate and digested overnight with 500ng/ul of trypsin (V5111, Promega) at 37°C. Digests were reduced, alkylated and desalted as described above.

All the samples (1ug digested peptides) were analyzed in triplicate by LC-MS/MS using a Q-Exactive mass spectrometer (Thermo Scientific, San Jose, CA) with the following conditions. A fused silica microcapillary column (100 mmID, 20 cm) packed with C18 reverse-phase resin (XSELECT CSH 130 C18 2.5 mm, Waters Co., Wilford, MA) using an in-line nano-flow EASY-nLC 1000 UHPLC (Thermo Scientific) was used to resolve the peptides. Peptides were eluted over a 45 min 2%–30% ACN gradient, a 5 min 30%– 60% ACN gradient, a 2 min 60%–95% gradient, with a final 8 min isocratic step at 0% ACN for a total run time of 60 min at a flow rate of 250 nl/min. All gradient mobile phases contained 0.1% formic acid. MS/MS data were collected in a data dependent fashion using a top 10 method with a full MS mass range from 300–1750 m/z, 70,000 resolution, and an AGC target of 3e6. MS2 scans were triggered when an ion intensity threshold of 1e5 was reached with a maximum injection time of 60 ms. Peptides were fragmented using a normalized collision energy setting of 25. A dynamic exclusion time of 20 s was used, and the peptide match setting was disabled. Singly charged ions, charge states above 8 and unassigned charge states were excluded.

The resultant RAW files were analyzed using Andromeda/MaxQuant (version 1.6.12.0) using the combined UniProt reviewed only database for Homo sapiens (Dec 2020). The default parameters were used and ‘match between the runs’ and ‘requantify’ options were enabled in the MaxQuant settings. The proteingroups output table was imported into Microsoft Excel for subsequent data analysis. Normalized SILAC ratios and LFQ intensities were used for data analysis.

### Purification of RNF10

Cells were seeded at 50% confluency in ten 10cm plates one day prior to transfection of a N-Flag-TEV-RNF10 expression plasmid using the calcium phosphate method. 20ug of total DNA was mixed with 2M CaCl2 in distilled water. The mixture was added in a dropwise manner to equal volumes 2XHBS (280mM NaCl, 10mM KCl, 1.5mM Na2HPO4, 12mM glucose and 50mM HEPES pH 7.05) solution at room temperature with continuous mixing, followed by incubation at room temperature for 30 minutes. Transfection mixture was added to each plate and incubated overnight at 37°C. 48 hours post transfection cells were collected by scrapping into cold 1X PBS and pelleted at 1,000 rpm for 5min at 4°C. Cells were lysed in 2mL of lysis buffer (50mM HEPES, pH 7.4, 100mM KAc, 5mM MgAc2, 0.5% NP40, 1 mM DTT (made fresh) and 1X EDTA-free Complete protease inhibitor cocktail) and incubated on ice for 20min. Lysates were clarified by centrifugation at 15,000 rpm for 10min at 4°C. 200ul of clarified lysate was added to a 1:1 slurry of pre-equilibrated (in lysis buffer with 0.1% NP40) anti-Flag M2 resin (A2220, Sigma) and incubated with rotation for 2 hours at 4°C. Resin was collected by centrifugation at 3,000 rpm for 1min at 4°C, while flow through was saved in a new tube. Resin was washed three times in 1ml of IP buffer (50mM HEPES, pH 7.4, 100mM KAc, 5mM MgAc2, 0.1% NP40, 1mM DTT (made fresh) and 1X EDTA-free Complete protease inhibitor cocktail) for 2min with rotation, followed by centrifugation. Resin was then washed three times with 1ml of high salt buffer (50mM HEPES, pH 7.4, 400mM KAc, 5mM MgAc2, 0.1% NP40, 1mM DTT), followed by three washes with 1ml of elution buffer (50mM HEPES, pH 7.4, 100mM KAc, 5mM MgAc2, 1mM DTT). Following elution, 100U of His-TEV protease (Z03030-1K, GenScript) was added to the 1:1 slurry of resin in elution buffer and incubated at room temperature for 30min. Resin was washed with an additional 100ul of elution buffer and then pooled with the first elution. 50ul of pre-equilibrated NiNTA agarose resin (30210, Qiagen) was added to the pooled elution fractions and incubated with rotation for 1h at 4°C. Cleared elution was collected by centrifugation, followed by silver stain and immunoblotting for confirmation of protein purification.

### In vitro ubiquitylation assay

All in vitro ubiquitylation reactions were carried out for 60min at 37°C. Single reactions consisted of 400nM recombinant human His6-Ubiquitin E1 enzyme Ube1 (E-304, BostonBiochem), 2uM recombinant human UbcH5c/UBE2D3 protein (E2-627, BostonBiochem), 200uM recombinant human ubiquitin no K (UM-NOK, BostonBiochem), 125nM 40S ribosomes (Purified from Hap1 cells, gift from Jody Puglisi and Alex Johnson, Stanford University), 50mM Tris-Cl pH 7.5, 50mM MgCl2, 20mM ATP, 6U/ml pyrophosphatase, 35U/ml creatine kinase and 100mM creatine phosphate, and 8uM RNF10. Reactions were inactivated with Laemmli buffer, then incubated for 10min at 95°C. Proteins were resolved by 12% SDS-PAGE and visualized by immunoblotting.

### Generation of knockout and knockin cell lines

Using CRISPR/Cas9 genome engineering USP10 and RNF10 knockout was done in 293Flp-In and 293T cells. Three individual guide RNAs were designed for each gene using CHOPCHOP website (https://chopchop.cbu.uib.no). RNF10: 5’-GCCGGCGAGTCTAAACCCAA-3’, 5’-GCCACGTTAGACTCGGGAAG-3’, 5’-CCGTTGATGCCGCTGAGCTC-3’, USP10: 5’-GACTCCTCGATCTTCAGTTG-3’, 5’-CTTACCTCAACTGAAGATCG-3’ and 5’-GCCTGGGTACTGGCAGTCGA-3. Cells were transfected with the pSpCas9(BB)-2a-GFP plasmid containing individual guide RNAs using lipofectamine 2000. 48 hours post transfection, GFP positive cells were either single cell sorted on a BD FACSAria Fusion (BD BioSciences) cell sorter, or pooled cell sorts were clonally isolated by limiting dilution method. Cells were validated for loss of USP10 and RNF10 by immunoblotting and sequencing. For HaloTag7 knock-in, guide RNA (gRNA) targeting the C-terminal region of human RPL26 gene was designed using the CHOPCHOP website (http://chopchop.cbu.uib.no/). The guide sequence for RPL26 gene (5′-GAAACCATTGAGAAGATGC-3′) was assembled into a pX459 plasmid. Donor vector was constructed by assembling a HaloTag7 transgene with upstream and downstream homology arms (650 nucleotide each) into a digested pSMART plasmid by Gibson assembly. Wild type HCT116 cells were transfected with donor and gRNA vectors (1 to 1 ratio) by Lipofectamine 3000 (Invitrogen). Five days after the transfection, the pool of transfected cells was treated with 100 nM Halo-TMR ligand for 1h, followed by washing three times. Fluorescence-positive cells were sorted into 96-well plates by flow cytometry (MoFlo Astrios EQ, Beckman Coulter). Three weeks later, the expanded single-cell colonies were screened for the integration of the HaloTag7 transgene by immunoblotting with α-RPL26.

### Ribo-Halo microscopy

HCT116 Ribo-HaloTag7 cells were transfected with either GFP-RNF10 WT or CS expression plasmid (2 ug/dish) using lipofectamine 3000 (Invitrogen). 24 hours post transfection, the cells were plated onto 35 mm-glass bottom dishes (No. 1.5, 14 mm glass diameter, MatTek) pre-treated with poly-L-lysine. 48 hours later, Halo-TMR containing medium (50 nM) was added to the cells and incubated for 1 hour. The medium was removed, and the cells were washed with warm DMEM for two times. DMEM was replaced by FluoroBrite™DMEM (Thermo Fisher) before the live cell imaging. The cells were imaged using a Yokogawa CSU-X1 spinning disk confocal with Spectral Applied Research Aurora Borealis modification on a Nikon Ti motorized microscope equipped with a Nikon Plan Apo 60×/1.40 N.A objective lens. Pairs of images for TMR and GFP fluorescence were collected sequentially using 100 mW 488 nm and 100 mW 561 solid state lasers attenuated and controlled with an AOTF (Spectral Applied Research LMM-5), and emission collected with a 525/50 nm or 620/60 nm filter (Chroma Technologies), respectively. Confocal images were acquired with the Hamamatsu ORCA-ER cooled CCD camera and MetaMorph software. The images were analyzed using FiJi software.

### Flowcytometry analysis for Ribo-Halo labeling

Ribo-Halo cells were seeded at 40% confluency in 12-well plates one day prior to transient transfections. 36 hours post transfection cells were treated with 100nM TMR-ligand (G8251, Promega) for 1-2 hours. After TMR-labeling, cells were washed with fresh warmed DMEM without the Halo-ligand three times with 10min incubations in between washes. Fresh warm DMEM was added to cells and cells were collected at various time points post washout. Cells were trypsinized then collected in fresh media. Following a short 3min centrifugation at 3,500rpm, cell pellets were resuspended in 800ul of FACS buffer (2% FBS in 1x DPBS) and passed through a nitex nylon mesh (Genesee Scientific). Samples were analyzed by flow-cytometry on a BD LSRFortessaTM X-20 cell analyzer (BD Biosciences). FACS data was analyzed using FlowJo (v10.6.2).

### qPCR analysis

For qPCR analysis, cells were plated at 50-60% confluency prior to lipofectamine based transfection, as described previously. 48 hours post transfection cells were collected in TRIzol and RNA was isolated using Direct-zol RNA miniprep kit (11-331, Zymo Research). Using 2ug RNA template, cDNA was synthesized is SuperScript III First Strand Synthesis system (18080-051, Invitrogen). Five standards were prepared by making four-fold dilutions of a sample pool. cDNA samples were each diluted 1:5 in water prior to plating. 8ul of each standard or sample was plated into a 96-well thermocycler plate, followed by 12ul of master mix containing SYBR green super mix (1725121, BioRad) and primers for gene of interest. The following primers were used in this study: RPS3: 5’-CAGAACAGAAGGGTGGGAAG-3’, 5’-GCAACATCCAGACTCCAGAATA-3’, RPS6: 5’-GAGCGTTCTCAACTTGGTTATTG-3’, 5’-GCGGATTCTGCTAGCTCTTT-3’, RPL7: 5’-GGCGAGGATGGCAAGAAA-3’, 5’-CTTTGGGCTCACTCCATTGATA-3’, GAPDH: 5’-AACGGGAAGCTTGTCATCAATGGAAA-3’, 5’-GCAGGAGGCAGCTGATGATCTT-3’. The following PCR conditions were run on a C1000 Thermo Cycler (BioRad): 50°C for 10min, 95°C for 15min, 95°C for 10s, 60°C for 30s (repeat for 40 cycles). All relative quantifications were calculated using the delta delta Ct method.

## QUANTIFICATION AND STATISTICAL ANALYSIS

All FACS-based assays were performed in triplicate (n = 3) as biologically distinct samples. The median 561nm/488nm ratio and SD were calculated. Transient overexpression experiments were compared to a transfection control. Immunoblot quantification of the relative % ubiquitylation and % phosphorylation was calculated by normalization of the individual intensities for each concentration to that of the no treatment control. Significance (p value) was calculated using an unpaired two-tailed Student’s t test using GraphPad Prism 9.0.

### Table of RNAi oligonucleotides

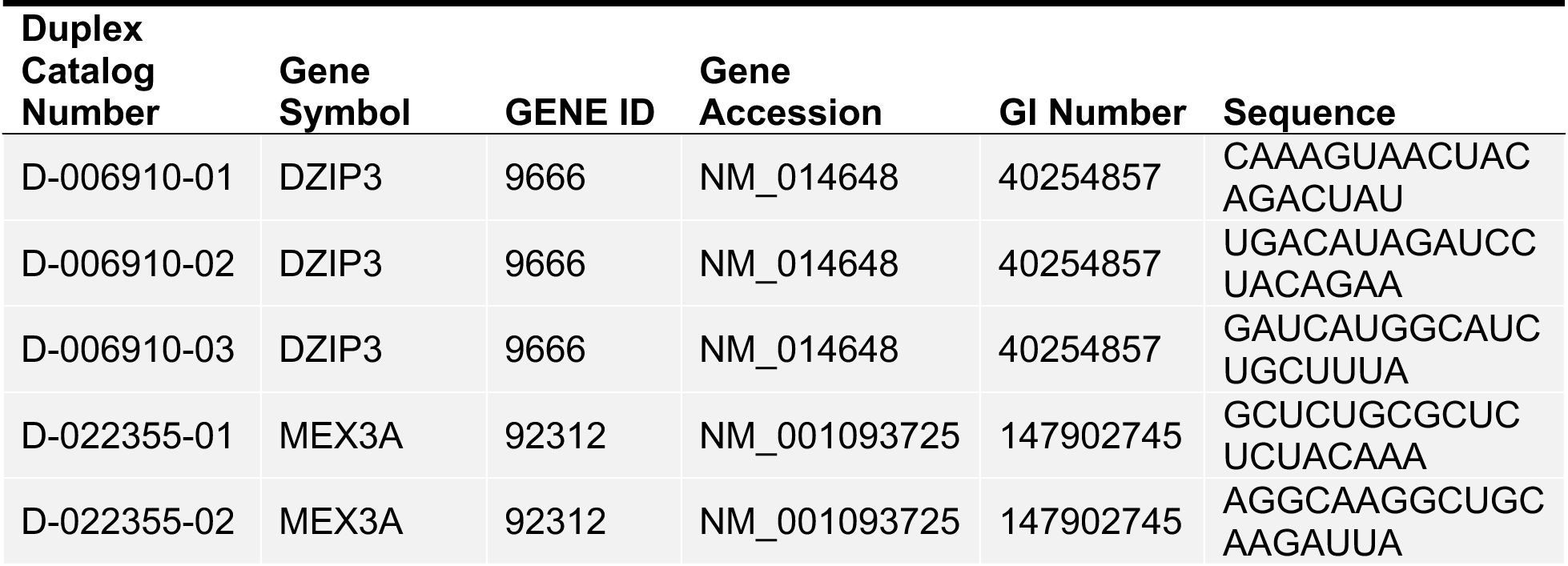
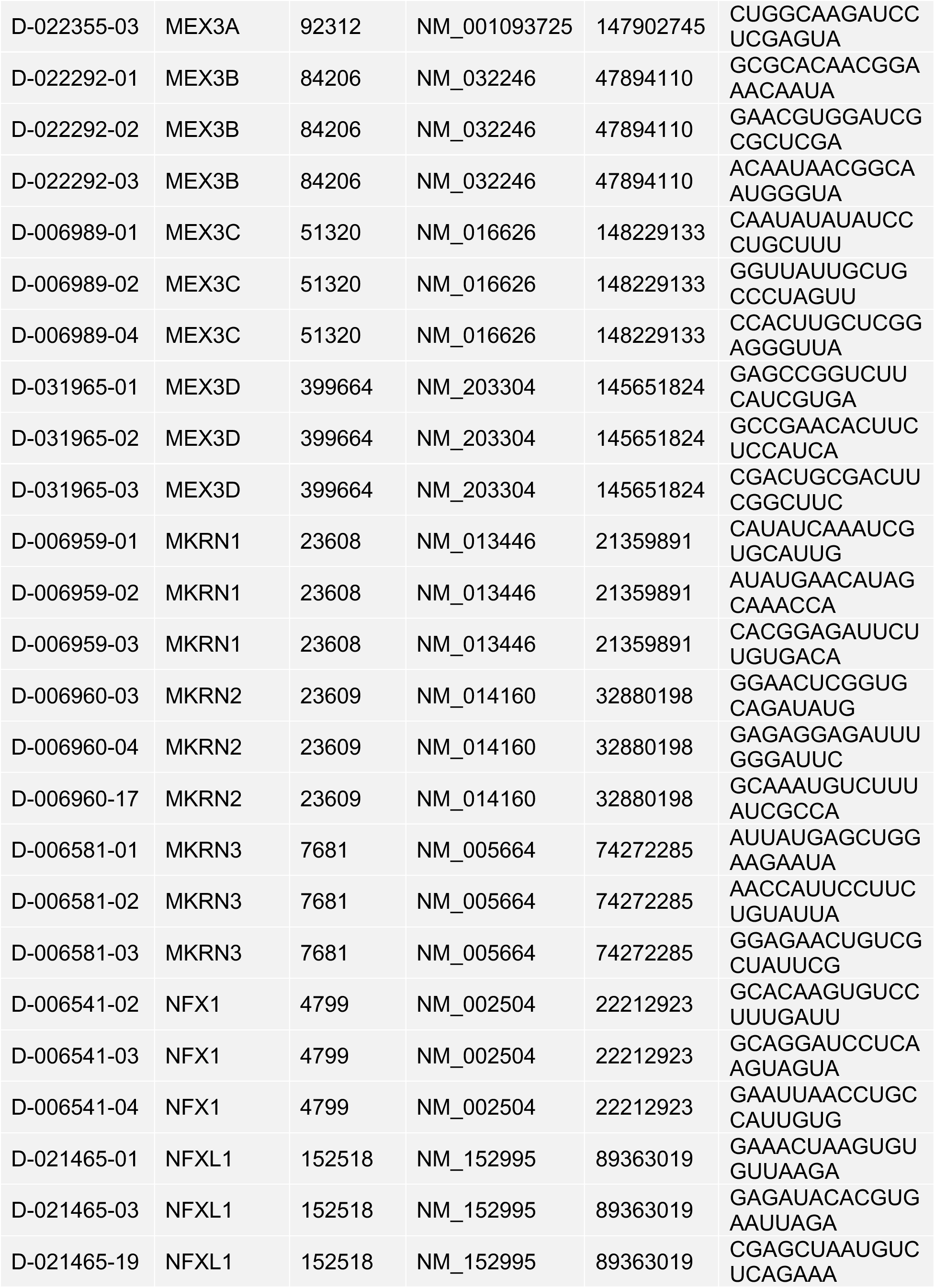
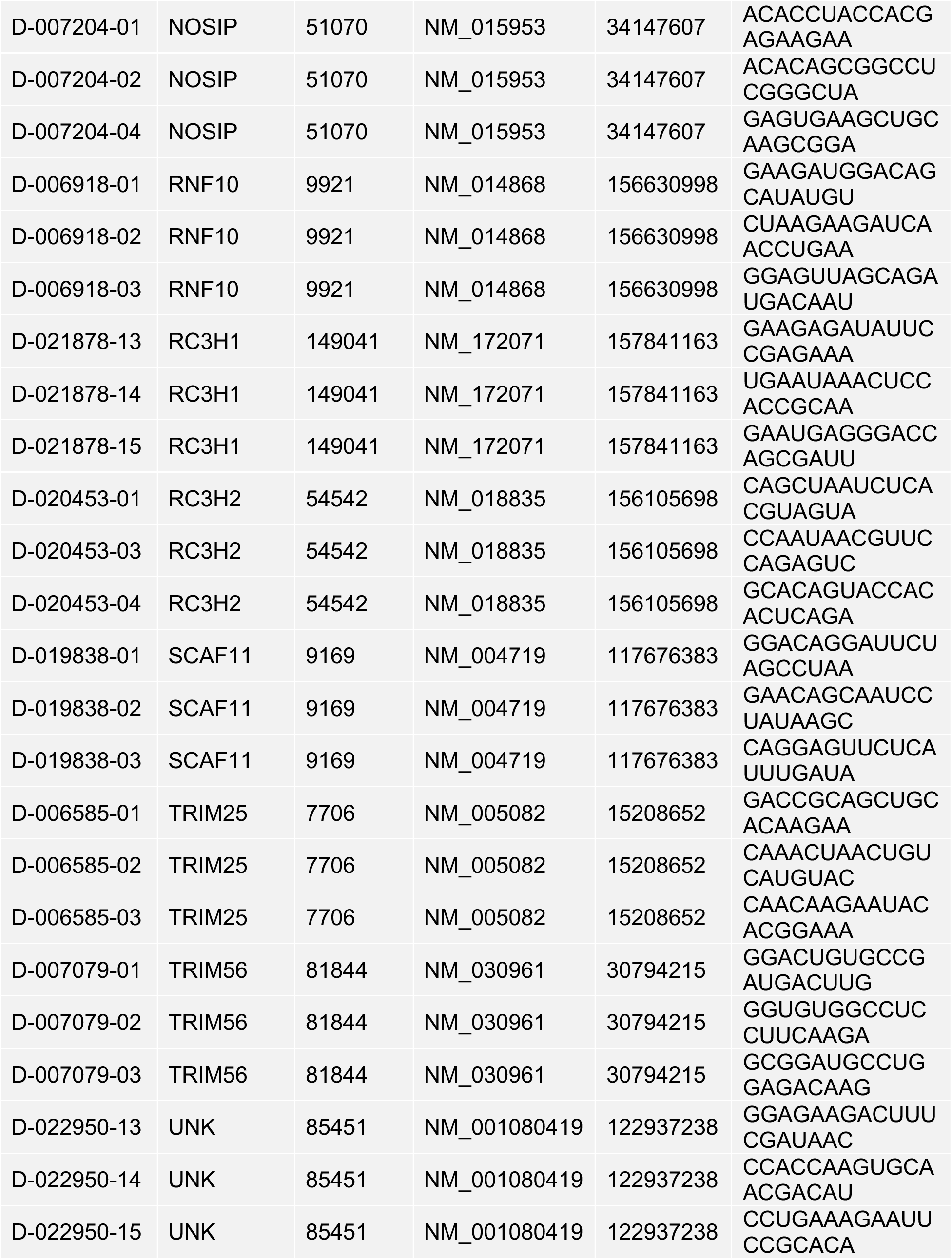
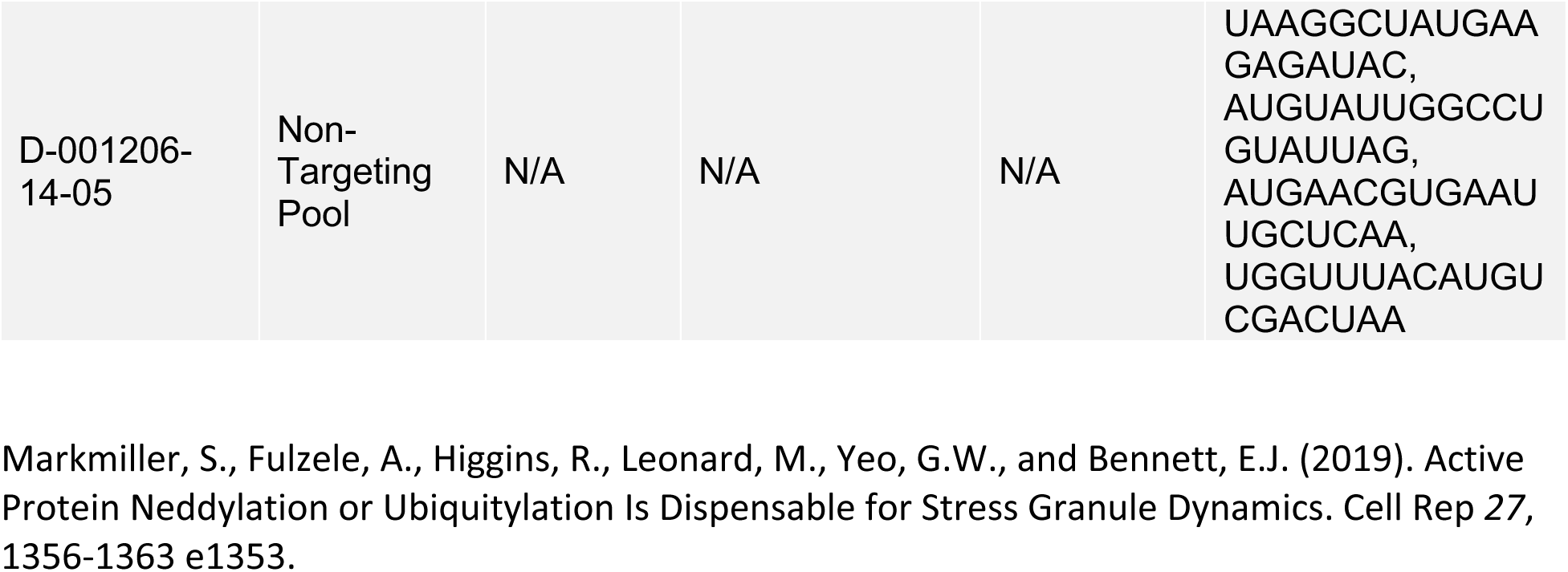

